# Time-resolved single-cell analysis of liver fibrosis resolution reveals a recovery-associated hepatic stellate cell state with prominent intercellular signaling

**DOI:** 10.64898/2026.11.25.769363

**Authors:** Eiko Saijou, Naoko Yokota, Tohru Itoh, Katsunori Fujiki, Katsuhiko Shirahige, Ryuichiro Nakato

## Abstract

**Background & Aims:** Activated hepatic stellate cells (HSCs) are central drivers of liver fibrosis, but their potential signaling roles during regression remain poorly understood. We investigated whether HSCs persisting after injury withdrawal express paracrine programs that could contribute to tissue recovery.

**Approach & Results:** We performed time-course single-cell RNA sequencing of enriched liver non-parenchymal cells during thioacetamide-induced fibrosis and after injury withdrawal in mice. During recovery, endothelial cells, macrophages, and cholangiocytes shifted toward healthy-like states, whereas HSCs retained a distinct transcriptional state. We found a persistent HSC state, which was abundant during both fibrosis and regression, showing declining apoptosis-related and activation-associated expression during recovery. We termed them recovery-associated HSCs (rHSCs). Intercellular communication analysis identified rHSCs as a prominent source of inferred outgoing signaling during early recovery. Gene regulatory network analysis nominated Wt1 and Mitf as candidate regulators of the associated ligand program, with Mdk emerging as a shared predicted target. We also detected an rHSC-like state in an independent mouse dataset, while human hepatic mesenchymal populations showed partial transcriptional correspondence. Combined delivery of five selected rHSC-associated factors reduced collagen deposition and increased the liver-to-body-weight ratio, serum total protein, and albumin.

**Conclusions:** These findings suggest that HSCs persisting during fibrosis regression may serve as signaling sources within the recovering liver rather than merely represent residual fibrogenic cells. The effects of selected factor delivery support the potential reparative relevance of the candidate paracrine program, while the contribution of endogenous rHSCs remains to be established. Animal study protocol approval number: A2022IQB014-05.

## INTRODUCTION

Liver fibrosis is a common consequence of chronic liver injury and is characterized by excessive deposition of extracellular matrix that progressively disrupts tissue architecture and function. (1) Although advanced fibrosis was once considered largely irreversible, clinical and experimental studies have established that fibrosis can regress after removal of the underlying injury. (2) Resolution involves degradation of scar matrix, remodeling of inflammatory and stromal cell populations, and restoration of tissue function. (3) However, the cellular programs that underlie this recovery process remain incompletely understood.

Hepatic stellate cells (HSCs) are major effector cells in liver fibrosis. In the healthy liver, quiescent HSCs (qHSCs) reside in the space of Disse and maintain a vitamin A-rich, non-fibrogenic state. (1) Following chronic injury, they acquire an activated myofibroblast-like phenotype and become a major source of fibrillar collagen and other extracellular-matrix components. (1) During fibrosis regression, activated HSCs can undergo apoptosis or transition toward less fibrogenic states, and lineage-tracing studies have shown that a subset survives injury withdrawal without fully returning to the original quiescent state. (4) Cellular senescence represents another potential fate of activated HSCs. Senescent HSCs have been reported to limit fibrosis and facilitate scar resolution, whereas therapeutic elimination of senescent cells can also ameliorate experimental liver fibrosis, suggesting context-dependent roles of HSCs for senescence in tissue injury and recovery. (5,6) These observations highlight substantial plasticity within the HSC compartment. However, whether persistent non-quiescent HSC populations simply represent residual fibrogenic cells or instead acquire active functions that contribute to tissue repair remains unresolved.

Fibrosis regression is a multicellular process involving coordinated responses among stellate cells, endothelial cells, macrophages, cholangiocytes, hepatocytes, and other hepatic populations. (2,3) These cells communicate through cytokines, growth factors, and other extracellular signals. (7) Single-cell transcriptomics provides an opportunity to examine cell-state dynamics together with inferred intercellular communication and gene-regulatory programs. However, time-resolved datasets spanning spontaneous fibrosis regression remain limited, particularly for relatively low-abundance non-parenchymal populations such as HSCs. (7–9)

Here, we investigated the cellular dynamics of liver non-parenchymal cells during thioacetamide (TAA)-induced fibrosis and spontaneous regression. To this end, we enriched liver non-parenchymal cells by fluorescence-activated cell sorting (FACS) and performed time-course single-cell RNA sequencing (scRNA-seq). The dataset captured established cell-type-specific transcriptional responses to liver injury and fibrosis. We identified a persistent HSC state with activation-associated features, termed recovery-associated HSCs (rHSCs), that was already abundant during fibrosis and remained prominent while undergoing transcriptional remodeling during regression. rHSCs were a prominent source of inferred outgoing signaling, and regulatory-network analysis nominated Wt1 and Mitf as candidate regulators of their ligand program. Combined delivery of five selected rHSC-associated candidate factors improved histological and serum biochemical readouts during fibrosis recovery. These findings link a persistent HSC state to a candidate paracrine program and suggest that fibrosis resolution may involve functional remodeling of activated HSCs rather than their simple disappearance or return to quiescence.

## MATERIALS AND METHODS

### Animals and experimental models of liver fibrosis and regression

All animal experiments were conducted in accordance with the Guidelines for the Care and Use of Laboratory Animals of The University of Tokyo and were approved by the Institutional Animal Care and Use Committee of the Institute for Quantitative Biosciences, The University of Tokyo (approval number A2022IQB014-05). Eight-week-old C57BL/6 mice were purchased from CLEA Japan (Tokyo, Japan) and maintained on standard chow (CLEA Rodent Diet CE-2 for mice, rats, and hamsters; CLEA Japan, Inc.) under a 12-h light/12-h dark cycle with ad libitum access to food and water. Animals were allocated to treatment groups without formal randomization, and investigators were not blinded to treatment allocation.

For the primary fibrosis/regression experiment, female mice received TAA (300 mg/L) in drinking water for 8 weeks and were analyzed during fibrosis or after 1, 2, or 4 weeks on normal drinking water. For the adeno-associated virus (AAV) experiments, male mice received TAA at 300 mg/L for 12 weeks, followed by AAV administration and analysis 2 weeks later. For the experiment using the midkine (MDK) inhibitor iMDK, male mice received TAA at 300 mg/L for 8 weeks, followed by daily iMDK administration for 2 weeks (14 administrations). Isoflurane anesthesia was used for procedures requiring anesthesia in the AAV experiments. Blood was collected from the retro-orbital plexus under anesthesia before euthanasia by cervical dislocation. Livers were excised into ice-cold PBS; the largest lobe was embedded in OCT compound (Sakura Finetek Japan), snap-frozen, and stored at -80 °C. Body and liver weights were recorded at the endpoint.

### Bulk RNA sequencing and transcriptomic analysis

Total liver RNA was isolated with TRIzol reagent (15596-026; Invitrogen). Libraries were prepared using the NEBNext Ultra Directional RNA Library Prep Kit for Illumina (E7420) and NEBNext rRNA Depletion Kit (E6310; New England Biolabs). Approximately 4 x 10^7 paired-end reads per library (36 bp x 2; insert size, approximately 240 bp) were generated on a NextSeq 2000 system (Illumina). The RNA-seq data was processed with RumBall (10). Briefly, the reads were mapped to GRCm38 with STAR (v2.7.10b), normalized expression levels were estimated with RSEM (v1.3.3), and differential expression was analyzed with DESeq2 (v1.36.0) (11–13), using a false discovery rate (FDR) cutoff of <0.01. Principal component analysis (PCA) and visualization were performed in R (v4.0.5). Pre-ranked Gene Set Enrichment Analysis (GSEA) compared TAA8w with recovery week 4 using Hallmark and Reactome gene sets (14,15). Detailed ranking, identifier-conversion, and gene-set parameters are provided in the Supplementary Methods.

We downloaded the processed RNA-seq data from the Gene Expression Omnibus (GEO) under the accession number GSE261310 (16), which included two biological replicates per condition.

### Isolation of liver non-parenchymal cells and fluorescence-activated cell sorting

Liver single-cell suspensions were prepared by pronase and collagenase perfusion as described previously (17). After erythrocyte lysis, propidium iodide was used to exclude dead cells. Cholangiocytes, macrophages, and liver sinusoidal endothelial cells (LSECs) were identified using antibodies against EpCAM, F4/80, and PECAM1, respectively, whereas HSCs were identified by endogenous vitamin A autofluorescence. Antibody and reagent details are provided in Supplementary Table 1. Because macrophage, LSEC, and HSC gates contained overlapping events attributable to aggregation, mutually exclusive negative gates were applied among the F4/80-positive, PECAM1-positive, and vitamin A-autofluorescent populations. Cholangiocytes were sorted by EpCAM expression without cross-exclusion. Doublets were excluded using pulse height and width.

### Single-cell RNA sequencing and primary data processing

Libraries were prepared with the Chromium Next GEM Single Cell 3-prime platform, Chip G, Single Cell 3-prime Library and Gel Bead Kit v3.1, and Single Index Kit T Set A (PN-1000213; 10x Genomics), and sequenced on a NextSeq 2000 using 28 cycles for Read 1, 8 cycles for Index 1, and 91 cycles for Read 2. The reads were mapped to refdata-gex-GRCm39-2024-A based on GRCm39 and GENCODE vM33 with Cell Ranger count (v8.0.0), using default cell calling without --expect-cells. The four samples include healthy control, TAA8w, recovery 1 week, and recovery 4 weeks. Each sample was derived from one individual mouse, yielding one biological sample and one scRNA-seq library per condition; samples were not pooled across animals.

### Single-cell transcriptomic analysis and cell-type annotation

Downstream single-cell analyses were performed using the ShortCake computational environment (18). Filtered feature-barcode matrices were analyzed with Seurat (v5.0.1). (19) Cells were retained when they contained >200 and <6,000 detected genes, <30,000 RNA counts, and <10% mitochondrial transcripts. For the cell-level analysis, the four datasets were merged. The log-normalized expression levels were scaled and analyzed by PCA. LSECs, HSCs, macrophages, and cholangiocytes were annotated by Pecam1, Lrat, Adgre1, and Epcam expression, respectively. Contaminating hepatocyte- or lymphoid-like clusters were excluded. Each cell type was reanalyzed separately. Cell-type-specific clustering and enrichment parameters are provided in the Supplementary Methods. For Figure 2F, unnormalized gene-level UMI counts were extracted from the Seurat RNA assay counts layer for LSECs, macrophages, cholangiocytes, and HSCs and summed across cells within each cell type and condition (healthy, TAA8w, recovery 1 week, and recovery 4 weeks), yielding 16 pseudobulk-like cell-type-by-condition profiles. Genes shared across the four cell-type-specific aggregated tables were retained; nonnumeric columns and genes with zero variance across the 16 profiles were removed. PCA was performed in R using prcomp with scaling enabled (scale. = TRUE).

We estimated co-expression similarity across HSCs based on the co-dependency index (CDI) using EEISP v0.6.4 (20). CDI quantified significant gene co-occurrence by applying a binomial test.

### Characterization of the rHSC transcriptional state

HSC_1 cells showed a quiescent-like transcriptional profile. HSC_2 was designated as the recovery-associated HSC (rHSC) state to reflect its association with the recovery-phase signaling program. The proportion of rHSCs among HSC_1 cells and rHSCs was calculated for each condition. Differential expression between rHSCs and HSC_1 cell was assessed with Seurat FindMarkers using log-normalized RNA values, min.pct = 0.1, and logfc.threshold = 0.25. Expression of representative quiescent, activated/fibrotic, and rHSC-associated genes was assessed by DotPlot, and Wt1 and Mdk distributions were visualized with Nebulosa.(21) Apoptosis, senescence, senescence-associated secretory phenotype (SASP), and detailed Gene Ontology (GO) analyses are described in the Supplementary Methods.

### Cell-cell communication analysis with CellChat

Cell-cell communication was inferred with CellChat (v1.6.1) (22) separately in healthy, TAA8w, recovery 1-week, and recovery 4-week samples. Six populations were analyzed: cholangiocytes, rHSCs, HSC_1 cells, Kupffer cells, LSECs, and monocyte-derived macrophages. The mouse CellChat database was restricted to Secreted Signaling interactions. Communication probabilities were inferred and summarized at pathway level, and condition-specific objects were compared to identify changes in outgoing and incoming signaling. rHSC-derived ligands associated with recovery were identified by integrating differential expression with inferred ligand-receptor interactions and ranking ligands increased at recovery 1 or 4 weeks. Detailed function-level settings and thresholds are provided in the Supplementary Methods.

### Gene-regulatory network analysis with CellOracle

Gene-regulatory networks(GRNs) were inferred with CellOracle (v0.16.0) (23) using the same 3,167-HSC cells analyzed with CellChat, comprising 622 HSC_1 cells and 2,545 rHSCs across healthy, TAA8w, recovery 1-week, and recovery 4-week samples. The input comprised 4,996 genes, with no additional cell filtering or reclustering. A prior regulatory network was generated from the mouse single-cell assay for transposase-accessible chromatin using sequencing (scATAC-seq) atlas supplied with CellOracle. Condition-specific GRNs were integrated with rHSC-associated ligands identified by CellChat and increased during recovery. Transcription factors connected to these ligands were evaluated based on their expression dynamics, prioritizing Wt1 and Mitf as candidate regulators. Cellular co-expression of Wt1 and/or Mitf with rHSC-associated ligands was calculated from log-normalized HSC expression data, defining detection as log1p-normalized expression >0. Detailed smoothing, network-estimation, and edge-filtering parameters are provided in the Supplementary Methods.

### Independent validation of the rHSC state in GSE295293

We downloaded the mouse liver scRNA-seq dataset from GEO under the accession number GSE295293, which contained three control, three TAA-treated fibrotic, and three post-TAA regression samples. (24) Samples were processed, integrated, annotated with established lineage markers with Seurat, and HSCs were extracted and reanalyzed using the RNA assay. Query HSCs were mapped to the saved PCA and uniform manifold approximation and projection (UMAP) representations of the primary HSC reference, in which cluster 0 was rHSC and the remaining HSC cluster was classified as other HSC. Both HSC cluster identities and the binary rHSC versus other-HSC annotation were transferred. An rHSC signature was constructed from positive cluster 0 markers identified by the Wilcoxon rank-sum test with min.pct = 0.10, logfc.threshold = 0.25, and adjusted P <0.05 after excluding mitochondrial, ribosomal, hemoglobin, cell-cycle, stress-response, and lineage-contamination genes. Wt1 and Mdk were explicitly excluded from the signature and reserved for independent validation. The top 30 remaining genes ranked by fold change were used for module scoring. Concordance between transferred labels and signature scores was evaluated, and Wt1 and Mdk detection frequency and average expression were compared between mapped rHSC-like and other HSC populations. Detailed integration and mapping settings are provided in the Supplementary Methods.

### Recombinant AAV vectors and in vivo administration

Recombinant adeno-associated virus (rAAV) vectors were produced in human embryonic kidney 293 cells as previously described. (25) The GFP transgene was removed from pLSP1eGFP to generate the pLSP1 expression backbone, and Igf1, Vegfc, Ngf, and Hgf coding sequences were cloned into this backbone. Because cloning of native Mdk was unsuccessful, a synthetic coding sequence, artMdk, was generated using synonymous substitutions that preserved the MDK amino acid sequence and enabled transgene-specific quantification. Constructs were pseudotyped with AAV8 by polyethyleneimine-mediated transfection using p5E18-VD2/8 and pXX6-80, yielding rAAV-GFP, rAAV-Igf1, rAAV-artMdk, rAAV-Vegfc, rAAV-Ngf, and rAAV-Hgf.

After 12 weeks of TAA treatment, male mice received intravenous rAAV and were analyzed 2 weeks later. Control mice received 1 × 10^11 vector genomes (vg) of rAAV-GFP, whereas the five-factor mixture group received 2 × 10^10 vg each of rAAV-Igf1, rAAV-artMdk, rAAV-Vegfc, rAAV-Ngf, and rAAV-Hgf, for the same total dose of 1 × 10^11 vg per mouse. Each group contained seven mice (n = 7).

### Histological analysis

Liver cryosections (8 micrometers) were fixed overnight at 4 degrees C in Bouin solution and stained with Direct Red 80 as described previously (16). Images were acquired with a BZ-X810 microscope (Keyence), and Sirius Red-positive areas were quantified at x20 magnification using ImageJ (v1.54e) (26). Fifteen fields were analyzed per liver, and the mean positive area was calculated for each animal.

### Quantitative real-time PCR

Total liver RNA was isolated with TRIzol, and cDNA was synthesized using PrimeScript RT Master Mix (Takara Bio). Quantitative real-time polymerase chain reaction (qPCR) was performed on a LightCycler 96 system (Roche Diagnostics). Relative expression was calculated using the 2^-DDCt method with Actb as the reference gene. artMdk-specific primers distinguished vector-derived from endogenous Mdk expression. Primer sequences are listed in Supplementary Table 2.

### Serum biochemical analysis

Whole blood was clotted at room temperature for 30 min and centrifuged at 250 × g for 10 min. Serum was stored at -20 °C. Total protein and albumin were measured with SPOTCHEM II LIVER-2 reagent strips (Arkray; 77323) according to the manufacturer’s instructions.

### Statistical analysis

Data are presented as specified in the figure legends. For two-group comparisons in mouse experiments, P values were calculated using two-sided unpaired Student’s t-tests. Each mouse was treated as an independent biological replicate for animal-level measurements. Individual cells were treated as observational units in cell-level analyses and were not considered as independent biological replicates; cell-level P values were therefore not interpreted as animal-level inference. Exact sample sizes and graphical summaries are indicated in the corresponding figure legends.

### Data availability

The primary scRNA-seq dataset generated in this study has been deposited in GEO under accession number GSE349112. Bulk RNA-seq data were obtained from GEO accession GSE261310 (16); the present study analyzed healthy, TAA8w, and recovery 1-, 2-, and 4-week samples and excluded choline-supplemented samples. Public single-cell datasets used for validation were obtained from GEO accessions GSE295293, GSE136103, and GSE202379.

## RESULTS

### Liver fibrosis regresses after withdrawal of chronic TAA exposure

To characterize spontaneous fibrosis regression, female mice were treated with TAA for 8 weeks and examined during fibrosis or at 1, 2, and 4 weeks after TAA withdrawal (Fig. 1A). Expression of the fibrosis-associated genes Timp1, Acta2, Col1a1, and Col3a1 declined during recovery, accompanied by reduced Sirius Red staining (Fig. 1B,C). We performed bulk liver RNA-seq analysis and found that fibrotic and recovery-stage samples separated in PCA, and that fibrogenic and hepatocyte function-associated genes showed differential expression (Fig. 1D,E). Selected Reactome pathways related to extracellular matrix organization and TGF-β signaling showed enrichment predominantly toward fibrotic liver rather than recovery week 4 (Fig. 1F). None of these pathways reached FDR <0.05. Hallmark GSEA analysis showed enrichment of cell-cycle and proliferative programs in fibrotic liver and of coagulation, bile acid metabolism, xenobiotic metabolism, and peroxisomal programs at recovery week 4 (Supplementary Fig. 1). Based on these temporal changes, we selected recovery weeks 1 and 4 as representative early and late phases of fibrosis regression, respectively, for subsequent scRNA-seq analysis.

**Figure 1.**
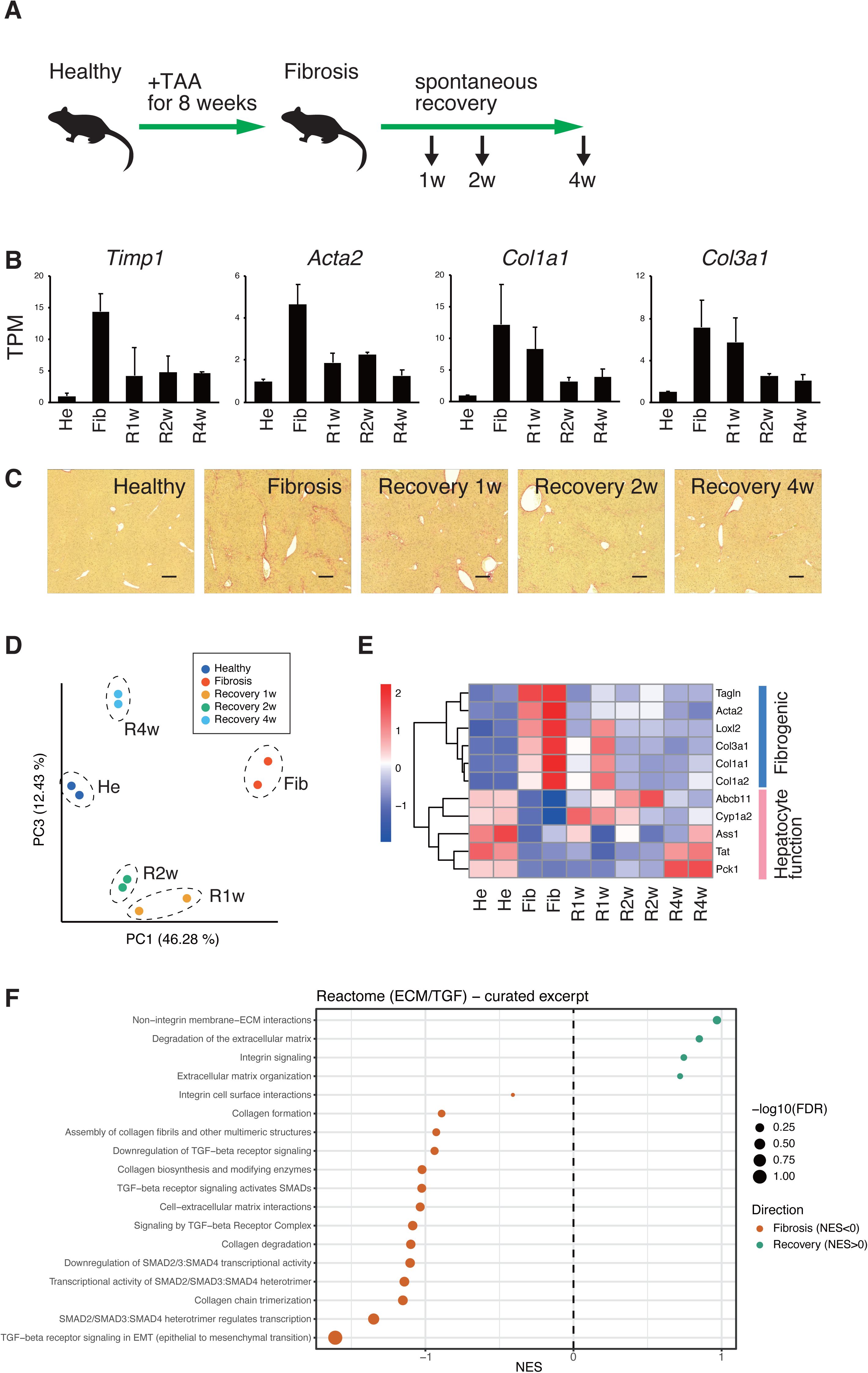
Transcriptional and histological changes during spontaneous regression of liver fibrosis. A, Schematic of the experimental model. Liver fibrosis was induced by thioacetamide (TAA) treatment for 8 weeks, followed by TAA withdrawal and spontaneous recovery for 1, 2, or 4 weeks. Healthy untreated mice were included as controls. B, Bulk RNA-seq expression levels of representative fibrosis-associated genes (Timp1, Acta2, Col1a1, and Col3a1) in healthy, fibrotic, and recovering livers at 1, 2, and 4 weeks after TAA withdrawal. Two biological replicates were analyzed per condition (n = 2). Bars show mean ± standard deviation (SD). C, Representative Sirius Red staining of liver sections from healthy mice, fibrotic mice after 8 weeks of TAA treatment, and mice after 1, 2, or 4 weeks of spontaneous recovery. Scale bar, 200 μm. D, PCA of bulk liver RNA-seq profiles from healthy, fibrotic, and recovery-stage samples. E, Heatmap showing expression of representative fibrogenic and hepatocyte function-associated genes across healthy, fibrotic, and recovery-stage livers. Expression values are shown as row-scaled values. F, Selected Reactome pathways related to extracellular matrix organization and TGF-β signaling from pre-ranked gene set enrichment analysis comparing fibrotic liver with recovery week 4. Negative normalized enrichment scores (NES) indicate enrichment on the fibrosis side of the comparison, and positive values indicate enrichment on the recovery side. Point size represents −log10(false discovery rate (FDR)). Pathways were selected for display based on their names without an FDR cutoff; displayed pathways did not reach FDR < 0.05.

### Single-cell profiling reveals distinct recovery dynamics among liver non-parenchymal cell populations

To characterize non-parenchymal cell states during fibrosis regression, we isolated liver endothelial cells, macrophages, HSCs, and cholangiocytes by FACS using PECAM1, F4/80, vitamin A autofluorescence, and EpCAM, respectively, and subjected the enriched populations to scRNA-seq (Fig. 2A). The resulting scRNA-seq data comprised 30,451 cells from healthy, fibrotic, recovery 1-week, and recovery 4-week livers. UMAP visualization and cell-type-associated marker expression confirmed representation of the four major non-parenchymal cell types (Fig. 2B–D).

**Figure 2.**
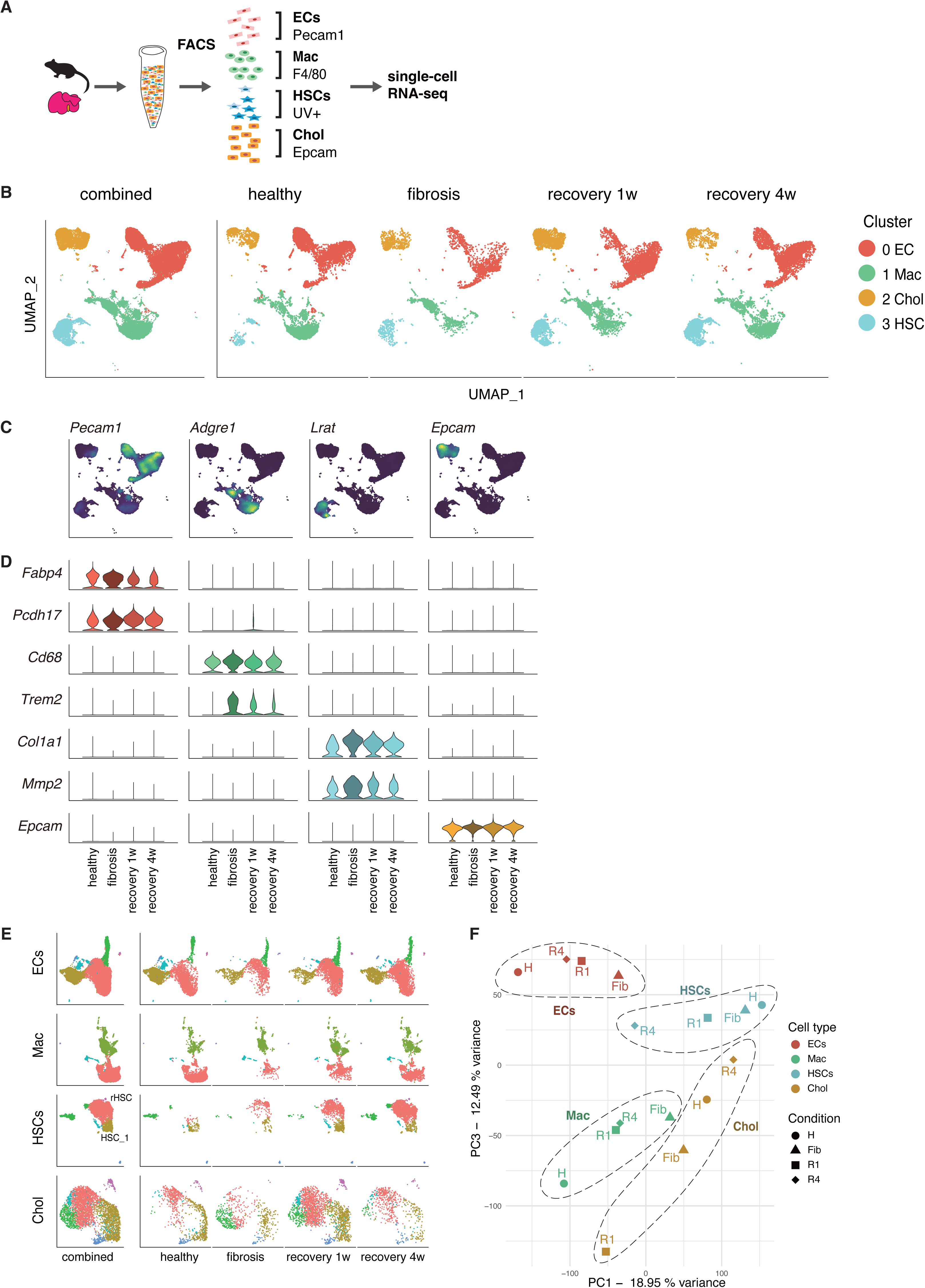
Single-cell RNA sequencing reveals cell-type-specific transcriptional remodeling during fibrosis regression. A, Schematic of the single-cell RNA-seq (scRNA-seq) workflow. Liver non-parenchymal cells were isolated by fluorescence-activated cell sorting (FACS), including endothelial cells (ECs; Pecam1^+^), macrophages (Mac; F4/80^+^), hepatic stellate cells (HSCs; vitamin A autofluorescence^+^), and cholangiocytes (Chol; Epcam^+^), followed by scRNA-seq. B, UMAP representation of the combined single-cell dataset and datasets split by experimental condition (healthy, fibrosis, recovery 1 week, and recovery 4 weeks). Cells are colored according to the four major cell populations: ECs, macrophages, cholangiocytes, and HSCs. Each condition was represented by one unpooled biological sample from one mouse (one scRNA-seq library). C, Density plots showing expression of representative lineage markers Pecam1, Adgre1, Lrat, and Epcam across the combined dataset, supporting the assignment of ECs, macrophages, HSCs, and cholangiocytes, respectively. D, Violin plots showing expression of representative cell-type- and state-associated genes across healthy, fibrotic, recovery 1-week, and recovery 4-week samples within each major cell population. E, UMAP representations of ECs, macrophages, HSCs, and cholangiocytes analyzed separately, shown as combined datasets and split by experimental condition, illustrating cell-type-specific changes in cellular states during fibrosis and regression. In the HSC panels, HSC_1 denotes cells with a quiescent-like transcriptional profile, and HSC_2 corresponds to recovery-associated HSCs (rHSCs). F, PCA of 16 pseudobulk-like profiles generated by summing raw counts across cells within each of the four major non-parenchymal cell types and each condition: healthy (H), fibrotic (Fib), recovery 1 week (R1), and recovery 4 weeks (R4). Each point represents one cell-type-by-condition aggregated profile. Colors indicate cell type (EC, macrophage, HSC, or cholangiocyte), and point shapes indicate experimental condition.

Compared with healthy samples, fibrotic samples showed increased expression of Col1a1 and Mmp2 in HSCs and Trem2 in macrophages. Endothelial cells also showed increased expression of Fabp4 and Pcdh17 (Fig. 2D). These patterns were broadly consistent with previously reported fibrogenic and matrix-remodeling responses, scar-associated macrophage features, and endothelial injury responses (27). Together, these observations indicated that the dataset captured established features of liver injury and fibrosis.

We next compared cell-state changes during recovery after injury withdrawal by subclustering each cell type separately (Fig. 2E). Endothelial cells, macrophages, and cholangiocytes showed distinct subcluster patterns during fibrosis, with recovery week 4 patterns more closely resembling those in healthy liver. In contrast, HSCs did not show a comparable return toward the healthy-state distribution. Within the HSC subclusters, HSC_1 predominated in healthy liver, whereas HSC_2 was abundant during fibrosis and remained prominent throughout recovery (Fig. 2E). PCA supported this contrast: recovery week 4 endothelial, macrophage, and cholangiocyte profiles shifted toward their corresponding healthy states, whereas the HSC profile remained separated from healthy HSCs (Fig. 2F). Endothelial cells showed increased expression of scavenger/endocytic receptor genes, including Stab2, Lyve1, and Cd36, at recovery week 1 relative to healthy liver (Supplementary Fig. 2A). Comparison with Kupffer cells identified Alcam, Sema4d, and Slamf7 as candidate surface-associated genes enriched in monocyte-derived macrophages (Supplementary Fig. 2B). In cholangiocytes, comparison of recovery week 4 with healthy liver identified changes in genes associated with apoptotic signaling and RNA regulation (Supplementary Fig. 2C). Together, these observations identified a persistent HSC state during fibrosis regression and prompted us to examine its potential role as a source of intercellular signals. We termed the HSC_2 subcluster recovery-associated HSCs (rHSCs), a designation that reflects their persistence during recovery rather than their exclusive presence at this stage (Fig. 2E; Supplementary Fig. 3A). Additional characterization showed declining apoptosis-related and activation-associated expression within rHSCs during recovery, consistent with transcriptional remodeling of this persistent state (Supplementary Fig. 3E). We focused our subsequent analyses on rHSCs.

### rHSCs are a prominent source of inferred intercellular signaling during early recovery

We inferred intercellular communication among cholangiocytes, HSC_1 cells, rHSCs, Kupffer cells, LSECs, and monocyte-derived macrophages with CellChat (Fig. 3). At recovery week 1, cholangiocytes showed the highest overall outgoing interaction strength in the complete network, largely reflecting persistent inferred SPP1-dominated signaling. Cholangiocytes were omitted from the sender-strength scatterplot to facilitate comparison among the other populations; among those displayed, rHSCs were a prominent source of inferred outgoing signaling (Fig. 3B). Differential analysis showed a marked increase in inferred outgoing MK signaling at recovery week 1 relative to healthy liver (Fig. 3C). In the inferred MK pathway network, healthy liver showed only weak cholangiocyte-derived signaling, whereas rHSCs emerged as a prominent predicted source at recovery week 1; no corresponding inferred MK output was detected from HSC_1 (Fig. 3D). Predicted ligand–receptor interactions connected rHSCs with multiple neighboring populations (Fig. 3E). rHSC-derived interactions meeting the recovery week 1 criterion included Mdk, Igf1, Vegfc, and Ngf, whereas Hgf did not meet the applied cutoff (Fig. 3F). At recovery week 4, Hgf, Igf1, Mdk, and Vegfc met the corresponding criterion, whereas Ngf did not (Supplementary Fig. 4F). Focused analysis further predicted persistent bidirectional communication between HSCs and LSECs across fibrosis and recovery, involving distinct ligand–receptor pathways in each direction (Supplementary Fig. 4G–I). Together, these analyses identified rHSCs as prominent predicted signaling senders during fibrosis regression, with outgoing ligand programs that differed between early and late recovery. Figure 3G summarizes these inferred signaling changes together with the transcriptional remodeling of rHSCs during recovery.

**Figure 3.**
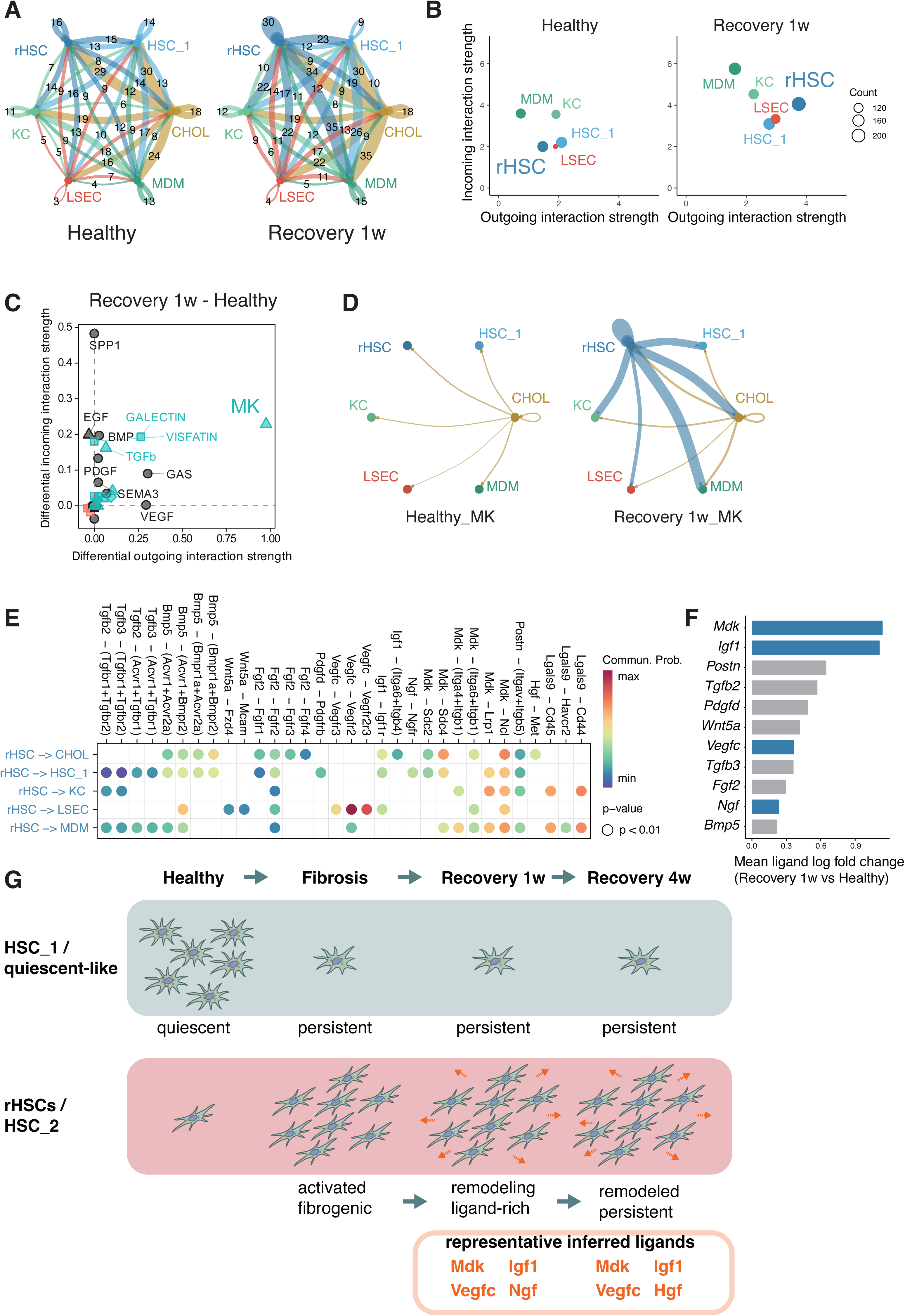
rHSCs are a prominent source of inferred intercellular signaling during early fibrosis regression. A, CellChat analysis of inferred intercellular communication networks in healthy liver and at 1 week of recovery after TAA withdrawal. Circle plots summarize predicted interactions among cholangiocytes (CHOL), HSC_1, rHSCs, Kupffer cells (KC), liver sinusoidal endothelial cells (LSEC), and monocyte-derived macrophages (MDM). Edge labels indicate the number of inferred ligand–receptor interactions. B, Comparison of outgoing and incoming interaction strengths among the displayed cell populations in healthy liver and at recovery 1 week. Cholangiocytes, which showed the highest overall outgoing interaction strength in the complete network, were omitted from this scatterplot because their persistently high SPP1-dominated inferred outgoing signal obscured comparison among the other populations. rHSCs were a prominent source of inferred outgoing signaling among the populations shown during recovery. C, Differential CellChat analysis comparing recovery 1 week with healthy liver. Signaling pathways are positioned according to changes in inferred outgoing and incoming interaction strength, highlighting pathways with increased inferred HSC-associated signaling during recovery, including MK, TGF-β, VEGF, SEMA3, GAS, BMP, and SPP1 signaling. MK is the CellChat pathway label for midkine (MDK) signaling. D, CellChat-inferred MK signaling networks in healthy liver and at recovery 1 week, showing changes in the predicted cellular sources and targets of MK-mediated communication. E, Inferred ligand–receptor interactions from rHSCs to cholangiocytes, HSC_1, KC, LSEC, and MDM at recovery week 1. Candidate pairs were selected by differentially expressed gene (DEG) mapping across all analyzed cell populations, rather than by ligand upregulation in rHSCs alone, and then displayed for rHSCs. Dot color represents inferred communication probability. F, Quantitative summary of the top rHSC-derived ligands showing increased inferred signaling at recovery 1 week (F1) relative to healthy liver. Ligands were ranked by mean ligand log fold change among F1-upregulated rHSC-derived CellChat interactions. Mdk, Igf1, Vegfc, and Ngf, four of the five factors subsequently used in the five-factor rHSC-associated mixture, are highlighted; Hgf did not meet the predefined F1-upregulation cutoff. G, Schematic summary of HSC states and inferred ligand programs during fibrosis regression. Quiescent-like HSC_1 cells coexist with HSC_2/rHSCs, which persist through recovery with reduced activation-associated expression and changing inferred ligand programs. Teal arrows indicate temporal progression and remodeling rather than lineage-traced transitions; orange dashed arrows indicate inferred outgoing signaling.

### CellOracle nominates Wt1 and Mitf as candidate regulators of the rHSC-associated ligand program

To investigate candidate regulatory relationships within the rHSC-associated ligand program, we used CellOracle, which infers gene regulatory networks using chromatin-accessibility-derived regulatory information together with scRNA-seq data. Using fifteen rHSC-associated candidate ligands identified by CellChat as the target set, CellOracle identified 33 candidate transcription factors with inferred connections to one or more ligands (Fig. 4A,B). We then examined their expression dynamics across healthy, fibrotic, and recovery-stage HSCs and prioritized Wt1 and Mitf as candidate regulators (Fig. 4C). CellOracle predicted connections from Wt1 and/or Mitf to multiple rHSC-associated ligands, including Mdk (Fig. 4D). For each condition, we quantified the proportion of HSCs in which each ligand was detected together with Wt1, Mitf, or both (Fig. 4E).

**Figure 4.**
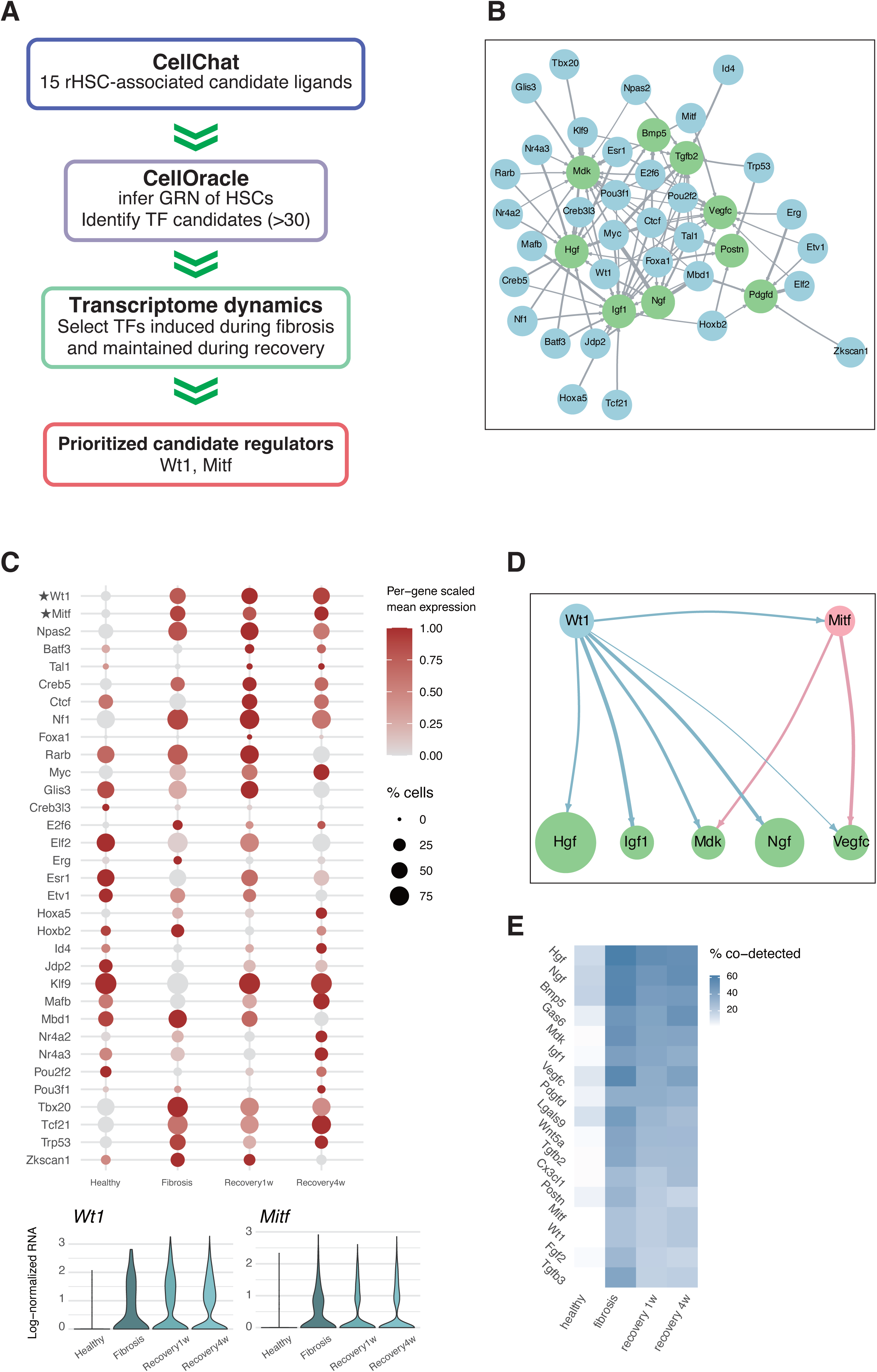
CellOracle analysis nominates Wt1 and Mitf as candidate regulators of the rHSC-associated ligand program. A, Schematic of the candidate-regulator nomination strategy. Candidate rHSC-associated ligands with increased inferred outgoing signaling at recovery 1 or 4 weeks, selected using the DEG-mapping and ligand fold-change criteria, were used as the target set for interrogation of the HSC gene regulatory network inferred by CellOracle. Candidate transcription factors with inferred connections to these ligands were nominated and further prioritized according to their expression dynamics across healthy, fibrotic, and recovery stages, yielding Wt1 and Mitf as prioritized candidate regulators. B, CellOracle-inferred gene regulatory network (GRN) linking candidate transcription factors to rHSC-associated ligands. Network structure was used to nominate transcription factors with predicted relationships to the rHSC-associated ligand program. Networks in B and D show recovery 1-week raw GRN edges without applying the P-value or top-2,000 edge filters used for target refinement in Supplementary Fig. 5A. C, Expression dynamics of CellOracle-nominated transcription factor candidates across healthy, fibrotic, recovery 1-week, and recovery 4-week conditions. Dot size indicates the percentage of cells expressing each transcription factor, and color indicates scaled expression. Violin plots below the dot plot show Wt1 and Mitf expression distributions across conditions. Wt1 and Mitf were prioritized as candidate regulators based on their expression dynamics during fibrosis and regression. D, CellOracle-inferred regulatory connections from Wt1 and/or Mitf to candidate rHSC-associated ligands showing increased inferred outgoing signaling at recovery 1 or 4 weeks based on the DEG-mapping and ligand fold-change criteria. All candidate target ligands with an inferred edge from either transcription factor are shown. E, Heatmap showing the proportion of HSCs co-expressing Wt1 and/or Mitf with rHSC-associated ligands across healthy, fibrotic, recovery 1-week, and recovery 4-week conditions. For each rHSC-associated ligand, the heatmap indicates the percentage of cells co-expressing that gene with either Wt1 or Mitf. For the Wt1 and Mitf entries, self-expression was excluded and co-expression with the other transcription factor was assessed. Percentages were calculated relative to all analyzed HSCs (HSC_1 cells and rHSCs) within each condition. Detection was defined as log-normalized RNA expression >0. Genes are ordered by decreasing co-expression frequency at recovery 1 week. Co-expression or co-detection does not establish direct transcriptional regulation.

We further refined the predicted Wt1 and Mitf targets using gene co-expression patterns across HSCs. This identified four candidate targets shared by Wt1 and Mitf at both recovery stages: Gas7, Plxna4, Runx1, and Serpine2 (Supplementary Fig. 5A–C). Together, these analyses prioritized Wt1 and Mitf as candidate regulators of the rHSC-associated ligand program, with Mdk among their predicted targets.

### External validation and cross-species evaluation of the rHSC-associated program

To determine whether the rHSC state was specific to our dataset, we examined the independent mouse dataset obtained from GSE295293. We applied reference mapping and identified an rHSC-like population comprising 1,167 of 4,590 HSCs (Supplementary Fig. 6E –G). This population showed higher rHSC signature scores and more frequent detection of Wt1 and Mdk, neither of which was included in the signature. rHSC-like cells were detected in both fibrogenic and regression samples (Supplementary Fig. 6H,I). These findings support the presence of an rHSC-like transcriptional state in an independent dataset, arguing against it being a dataset-specific observation. We also analyzed human datasets from GSE136103 (28) and GSE202379 (29) and observed partial correspondence of murine rHSC-associated transcriptional features with human hepatic mesenchymal populations, although no directly equivalent WT1+MDK+ human rHSC population was identified (Supplementary Fig. 7).

### Combined delivery of five rHSC-associated factors improves selected recovery-associated readouts in vivo

We then examined whether MDK alone could influence fibrosis regression. MDK overexpression significantly reduced hepatic Col3a1 expression, but did not significantly alter Acta2, Col1a1, Tgfb1, and Timp1 expression and Sirius Red-positive area. Pharmacological MDK inhibition with iMDK likewise produced no significant changes in the measured histological or transcriptional endpoints (Supplementary Fig. 8). Thus, manipulating MDK alone yielded limited effects under the conditions tested.

We next tested combined delivery of five rHSC-associated candidate factors highlighted by the signaling and regulatory analyses. Following 12 weeks of TAA treatment, male mice received either control rAAV-GFP or a mixture of rAAV vectors encoding IGF1, MDK (encoded by the synthetic artMdk sequence), VEGFC, NGF, and HGF; mice were assessed 2 weeks later. Hepatic expression of the delivered factors increased, whereas measured fibrosis-associated transcripts showed limited changes (Fig. 5A). Despite these limited transcriptional changes, the five-factor mixture significantly reduced Sirius Red-positive area compared with controls, demonstrating reduced collagen deposition (Fig. 5B, C). Mixture-treated mice also showed a higher liver-to-body-weight ratio without a significant difference in body weight (Fig. 5D). Importantly, serum total protein and albumin concentrations were both significantly increased, consistent with improved hepatic synthetic function (Fig. 5E). Together, these results demonstrate that combined delivery of rHSC-associated factors reduces collagen deposition and improves serum biochemical markers associated with hepatic synthetic function during fibrosis recovery.

**Figure 5.**
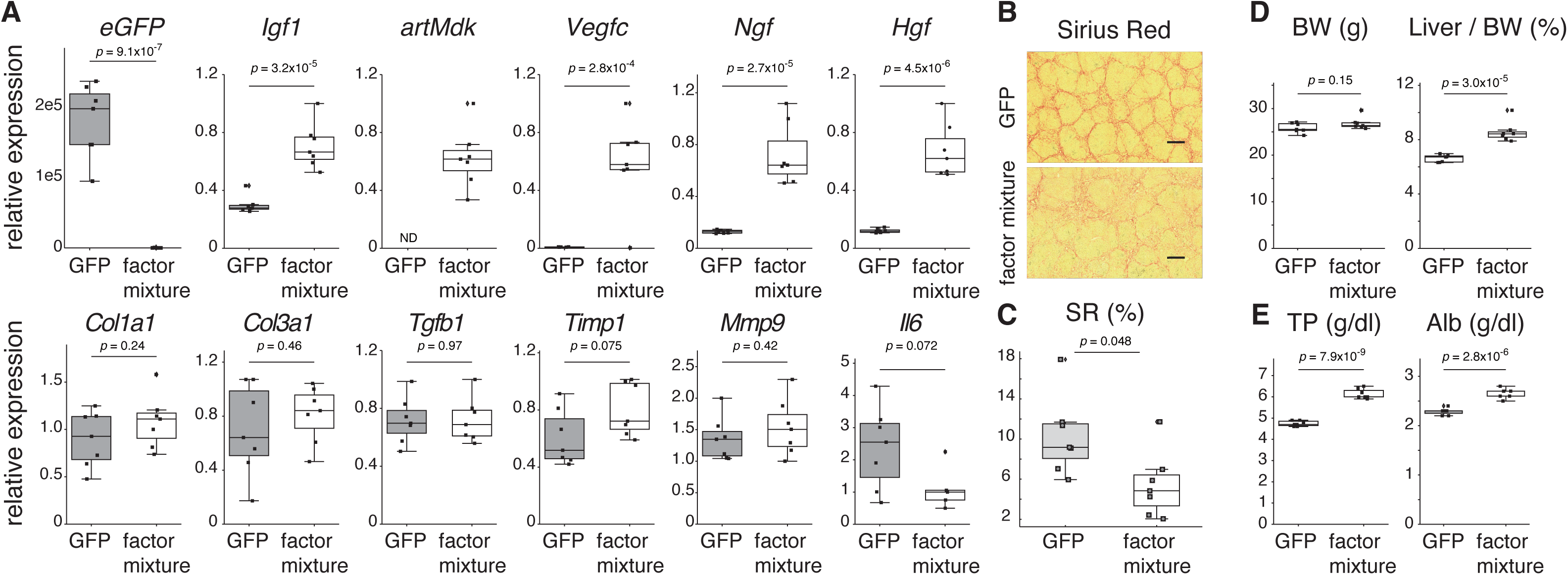
Combined delivery of rHSC-associated factors improves selected histological and serum biochemical measures during fibrosis recovery. Mice were treated with TAA for 12 weeks to establish liver fibrosis and then administered a mixture of recombinant adeno-associated virus (rAAV) vectors encoding five rHSC-associated factors (factor mixture; Igf1, artMdk, Vegfc, Ngf, and Hgf) or control rAAV-GFP. Mice were analyzed 2 weeks after rAAV administration (n = 7 mice per group). Box plots show the median and interquartile range (25th–75th percentiles), with whiskers extending to the most extreme values within 1.5 × the interquartile range; individual animals are shown as points. P values for qPCR (A), body weight and liver-to-body-weight ratio (D), and serum measurements (E) were calculated using two-sided Student’s t-tests. A, Hepatic expression of the five delivered rHSC-associated factors and fibrosis-associated genes in rAAV-GFP-treated and factor-mixture-treated mice. Expression of the delivered factors was increased in the factor-mixture group, whereas expression of fibrosis-associated genes showed limited overall changes. Statistical comparisons were not performed for transgene expression when expression was undetectable in one of the groups. B, Representative Sirius Red staining of liver sections from rAAV-GFP-treated and factor-mixture-treated mice. Scale bar, 200 μm. C, Quantification of Sirius Red-positive area, showing a lower Sirius Red-positive area in mice treated with the factor mixture. The P value for Sirius Red-positive area was calculated using a two-sided Student’s t-test. D, Body weight and liver-to-body-weight ratio 2 weeks after AAV administration. Factor-mixture treatment increased the liver-to-body-weight ratio without a significant change in body weight. E, Serum total protein (TP) and albumin (Alb) concentrations. Both TP and albumin were significantly increased in mice treated with the factor mixture. Exact P values are shown in the graphs.

## DISCUSSION

Fibrosis regression was accompanied by persistence and remodeling of an HSC state that remained transcriptionally distinct from healthy HSCs. This suggests that tissue recovery can coexist with non-quiescent stromal states. Previous studies have examined apoptosis and inactivation of activated HSCs during regression, as well as the role of HSC senescence in limiting fibrosis (4,6). Our findings extend this framework by suggesting that persistent non-quiescent HSC states may serve as signaling sources during tissue recovery. Differences between early and late inferred ligand programs suggest that the potential interactions of rHSCs with neighboring cells change during recovery. Predicted reciprocal communication with LSECs further raises the possibility that HSCs both respond to and influence the recovering tissue environment.

Previous single-cell atlas analyses have identified conserved transcriptional regulatory programs during HSC activation across mouse liver injury models, including Wt1-, Prrx1-, and Mef2c-associated regulon activity in myofibroblastic HSCs (30). The persistence of rHSCs during fibrosis regression, despite reduced fibrogenic gene expression, raises the possibility that these cells retain functions beyond the intercellular signaling programs inferred by CellChat. In particular, cell-intrinsic transcriptional programs involved in extracellular matrix remodeling warrant further investigation.

CellOracle analysis and expression dynamics prioritized Wt1 and Mitf as candidate regulators, with Mdk among their predicted targets. The identification of Wt1 is notable because injury-induced WT1 can restrain fibrogenesis. Kendall et al. showed that WT1 is induced in activated HSC populations, yet deleting Wt1 in PDGFRβ-expressing HSCs and myofibroblasts exacerbates fibrosis (31). Their study implicated Wnt signaling, matrix-associated programs, and myofibroblastic transition. Our findings raise an additional possibility: WT1 may coordinate a paracrine program with reparative potential in persistent HSC states. This hypothesis links the previously reported antifibrotic role of WT1 to candidate signaling functions during recovery.

Prior studies provide a regenerative context for MDK signaling. Loss of mdka in zebrafish reduced endothelial proliferation and resulted in persistence of a collagen-rich scar after cardiac injury (32). Activated HSC-derived Mdka promoted bipotential progenitor cell redifferentiation during zebrafish biliary-mediated liver regeneration (33). These observations provide a biological rationale for investigating MDK within an HSC-associated recovery program.

Combined delivery of five rHSC-associated factors reduced collagen deposition and increased serum total protein and albumin, biochemical measures associated with hepatic synthetic function. These changes provide experimental support for the reparative potential of selected components of the inferred ligand program. MDK overexpression alone had limited effects, and iMDK did not significantly alter measured endpoints under the conditions tested (Supplementary Fig. 8). Neither experiment establishes whether MDK is necessary for the mixture effect. HGF has recognized regenerative and antifibrotic activities (34). The individual contributions and any cooperative effects among the five factors remain unresolved. Hepatic factor delivery also does not establish that endogenous rHSCs produce the same effects.

Detection of an rHSC-like state in an independent mouse dataset supports recurrence of the transcriptional state beyond the primary dataset. Human hepatic mesenchymal populations showed partial transcriptional correspondence, but a directly equivalent WT1+MDK+ population was not identified. The available human datasets do not sample defined fibrosis-regression time courses. Species differences, disease context, and sampling therefore limit interpretation of the human correspondence.

Several limitations constrain these interpretations. Each condition in the primary single-cell time course was represented by one mouse, limiting assessment of between-animal variation. Individual cells cannot substitute for independent biological replicates. FACS enrichment improved representation of selected non-parenchymal populations but may bias the cell states recovered and does not provide a comprehensive liver atlas.

CellChat and CellOracle infer communication and regulatory relationships rather than directly measuring signaling activity or transcriptional regulation. Wt1- and Mitf-dependent regulation of the associated ligand program, including Mdk, remains to be tested. The temporal comparisons also do not establish the lineage origin of rHSCs or whether the same cells persist throughout recovery. Isolated-cell cultures may not reproduce the multicellular environment of the recovering liver. Constitutive knockout may confound effects on fibrosis development with those on regression. Perturbations lacking cell-type specificity may also affect other liver cell populations. Lineage tracing and temporally controlled, HSC-targeted perturbations during recovery will therefore be needed to test the endogenous contributions of rHSCs.

Despite these limitations, our findings suggest that persistent HSC states may participate in recovery-associated signaling and identify a factor combination with beneficial histological and serum biochemical effects. Determining whether and how endogenous HSCs use these candidate programs will clarify their contribution to fibrosis resolution and their potential for promoting tissue repair.

## Author contributions

E.S.: Conceptualization, Methodology, Investigation, Formal analysis, Data curation, Visualization, Project administration, Funding acquisition, Writing – original draft, Writing – review & editing. N.Y.: Investigation. T.I.: Conceptualization, Investigation, Resources, Funding acquisition, Writing – review & editing. K.F.: Resources. K.S.: Resources, Funding acquisition, Supervision. R.N.: Conceptualization, Methodology, Supervision, Funding acquisition, Writing – review & editing.

## Acknowledgments

We thank Yoshiko Kamiya for assistance with the experiments, Chizuko Koga for assistance with fluorescence-activated cell sorting, Yuya Nagaoka for technical assistance with iMDK administration, Testuo Oikawa for suggesting the in vivo factor-delivery experiment, and Taketomo Kido and Atsushi Miyajima for providing laboratory facilities and experimental space. OpenAI ChatGPT and Codex were used to assist with manuscript organization, English language editing, and the development of data-analysis and figure-generation code. The authors reviewed the AI-assisted work and take responsibility for the final content and interpretation.

## Abbreviations

AAV: adeno-associated virus;
ECM: extracellular matrix;
FACS: fluorescence-activated cell sorting;
HSC: hepatic stellate cell;
LSEC: liver sinusoidal endothelial cell;
MDK: midkine;
rAAV: recombinant adeno-associated virus;
rHSC: recovery-associated hepatic stellate cell;
scRNA-seq: single-cell RNA sequencing;
TAA: thioacetamide.

