## Supplemental text for "Time-resolved single-cell analysis of liver fibrosis resolution reveals a recovery-associated hepatic stellate cell state with prominent intercellular signaling"

### SUPPLEMENTARY METHODS

#### Detailed bulk RNA sequencing gene set enrichment analysis

Pre-ranked GSEA used the DESeq2 comparison between TAA8w and recovery week 4. Genes were ranked by the signed Wald statistic calculated from the log2 fold change and its standard error; positive values indicated higher expression during recovery and negative values indicated higher expression in fibrotic liver. Ensembl identifiers were converted to Entrez Gene identifiers with org.Mm.eg.db. When multiple entries mapped to one Entrez identifier, the entry with the largest absolute ranking statistic was retained.

Hallmark enrichment was performed with clusterProfiler (v4.12.6) using Hallmark sets obtained from MSigDB through msigdbr. Gene sets containing 10-2,000 genes were included. Reactome pre-ranked GSEA used gsePathway in ReactomePA (v1.48.0), with gene-set sizes restricted to 5-2,000 genes. Normalized enrichment scores indicated direction and magnitude; false discovery rates were calculated by the Benjamini-Hochberg method. Reactome pathways related to extracellular matrix organization and TGF-beta signaling supported Main Figure 1, whereas global Hallmark enrichment supported Supplementary Figure 1.

#### Lineage-specific single-cell analysis and functional enrichment

Seurat objects were created with min.cells = 3 and min.features = 200. After merging and log normalization, highly variable genes were identified with FindVariableFeatures, expression was scaled with ScaleData, and PCA was performed with RunPCA. The shared nearest-neighbor graph and UMAP used the first 30 principal components; initial graph-based clustering used resolution 0.03. HSCs, macrophages, LSECs, and cholangiocytes were then reanalyzed separately with 30 principal components. Clustering resolutions were 0.2, 0.03, 0.1, and 0.3, respectively. Cluster identities were assigned from canonical markers and cluster-enriched genes.

Differential expression used Seurat FindMarkers. For the non-HSC analyses, genes expressed in at least 25% of cells in either group were tested with a minimum log2 fold-change threshold of 0.25. Endothelial cells were compared between healthy and recovery 1-week samples, and cholangiocytes between healthy and recovery 4-week samples. Macrophage clusters were assigned as Kupffer cells using Clec4f and Vsig4 or monocyte-derived macrophages using Cx3cr1 and Trem2, and these populations were compared within each condition.

For GO analysis, genes with adjusted P <0.01 were divided into upregulated and downregulated sets by the sign of the average log2 fold change. Symbols were converted to Entrez identifiers with org.Mm.eg.db, and clusterProfiler::enrichGO was applied to Biological Process, Molecular Function, and Cellular Component ontologies with Benjamini-Hochberg correction, pvalueCutoff = 0.01, and qvalueCutoff = 0.05. These analyses supported Supplementary Figure 2.

**Detailed CellChat implementation**

For each condition, normalized expression and cell annotations were used to create a CellChat object. Ligand-receptor analysis used CellChatDB.mouse restricted to Secreted Signaling. Overexpressed genes and interactions were identified with identifyOverExpressedGenes and identifyOverExpressedInteractions. Expression was projected onto the mouse protein-protein interaction network with projectData. Communication probabilities were inferred with computeCommunProb; interactions involving groups with fewer than 10 cells were removed with filterCommunication. Probabilities were summarized with computeCommunProbPathway, networks were aggregated with aggregateNet, and pathway-level centrality was calculated with netAnalysis_computeCentrality.

Condition-specific objects were merged with mergeCellChat. Healthy liver was compared with recovery 1 and 4 weeks at pathway and ligand-receptor levels. Differentially regulated signaling was identified by mapping condition-specific differential expression onto the inferred CellChat interactions. For the recovery week 1 comparison, identifyOverExpressedGenes used group.dataset = "datasets", pos.dataset = "F1", only.pos = FALSE, thresh.pc = 0.1, thresh.fc = 0.1, and thresh.p = 1. The resulting features were mapped to inferred ligand–receptor interactions with netMappingDEG. Recovery week 1-associated interactions were then extracted with subsetCommunication using datasets = "F1" and requiring ligand log fold change >0.2, without a receptor fold-change threshold (receptor.logFC = NULL). No additional interaction-level P-value or multiple-testing threshold was applied at this DEG-mapping step. Recovery 1- and 4-week analyses were performed independently, and rHSC-derived ligands were ranked by mean ligand log fold change across retained interactions.

##### HSC-LSEC communication analysis

HSCs and LSECs were extracted from the integrated Seurat object and analyzed separately with CellChat. HSCs were treated as one population without subdivision into HSC_1 cells and rHSCs. Objects were generated independently for healthy, TAA8w, recovery 1-week, and recovery 4-week samples using the mouse Secreted Signaling database and the inference workflow described above. Communication probabilities were inferred with computeCommunProb, and interactions involving cell groups with fewer than 10 cells were removed using filterCommunication(min.cells = 10). The objects were merged with mergeCellChat; the number and strength of inferred interactions were compared across conditions, and HSC-to-LSEC, HSC-to-HSC, LSEC-to-HSC, and LSEC-to-LSEC interactions were displayed with netVisual_bubble using the native CellChat interaction P-value threshold of $P < 0.05$. This threshold applies to communication-probability inference and does not test differential signaling between conditions. This analysis supported Supplementary Figure 4.

**Detailed CellOracle implementation**

CellOracle (v0.16.0) analysis included the same 3,167 HSCs used in CellChat analysis, comprising 622 HSC_1 cells and 2,545 rHSCs, without additional cell filtering or reclustering. Of the original 4,999-gene list, 4,996 genes were present in the Seurat inputs and retained. Nonnegative integer counts from the Seurat RNA counts layer were imported using import_anndata_as_raw_count, with hsc_state as the cell-state annotation and the retained Seurat UMAP coordinates as X_umap. The saved TF-target prior dictionary from the original CellOracle analysis was reused.

Fifteen principal components were retained from CellOracle PCA. Expression was smoothed by k-nearest-neighbor imputation using 79 neighbors, approximately 2.5% of HSCs, with balanced = TRUE, b_sight = 632, b_maxl = 316, and n_jobs = 4. Condition-specific networks were inferred jointly from HSC_1 and rHSCs within each condition using oracle.get_links, with batch as the GRN unit, alpha = 10, model_method = 'bagging_ridge', bagging_number = 20, and n_jobs = 4. Edges were filtered at $P < 0.001$, ranked by absolute regression coefficient (coef_abs), and limited to 2,000 per condition.

rHSC-derived ligands with increased inferred signaling at recovery 1 or 4 weeks were used as targets. TFs with predicted edges to at least one target in either recovery network were combined into a nonredundant set. Candidate TFs were defined as the union of TFs connected to at least one of the 15 rHSC-associated ligands in the filtered recovery 1- or 4-week networks, yielding 33 candidates. Their expression profiles were then examined to prioritize factors induced during fibrosis relative to healthy HSCs and maintained during recovery, leading to the selection of Wt1 and Mitf.

For co-expression analysis, a gene was detected when log1p-normalized expression was >0. The proportion of HSCs co-expressing each rHSC-associated ligand with Wt1 or Mitf was calculated in each condition. For Wt1 and Mitf entries, self-expression was excluded. Frequencies were expressed as percentages and ordered by recovery 1-week frequency.

**CellOracle-CDI integration and NicheNet analysis**

Co-expression modules were identified using CDI as implemented in EEISP v0.6.4. The union of two modules containing both Wt1 and Mitf (modules 5 and 437; 368 unique genes) was integrated with the filtered CellOracle-predicted target sets. Wt1 and Mitf target sets were obtained separately for recovery weeks 1 and 4 using $P < 0.001$ and the top 2,000 edges by absolute coefficient per condition, and intersected with the CDI gene set. Genes shared by both transcription factors at both recovery stages were retained, yielding Gas7, Plxna4, Runx1, and Serpine2. Their observed log-normalized HSC expression was summarized across conditions. Recovery 1-week edges were displayed with width scaled to the absolute regression coefficient and color indicating the source TF; the additional Wt1-to-Mitf connection was displayed from the raw GRN.

NicheNet analysis used nichenetr (v2.2.1.1), with the same 3,167 HSCs analyzed with CellChat and CellOracle as receivers. LSECs, macrophages, cholangiocytes, and these HSCs were candidate senders. Receptor and background-gene filtering and sender-expression summaries pooled all four conditions; genes detected in at least 5% of cells within each analyzed population were considered expressed. Recovery 1-week versus TAA8w differential-expression results were generated separately and did not define the response gene set. The response set comprised all 33 candidate TFs identified by the updated CellOracle analysis (Fig. 4C). Standard mouse ligand-receptor, ligand-target, and weighted signaling priors were used. Candidate ligands required an expressed cognate receptor in HSCs; a sender-filtered set additionally required expression in at least one sender population. Ligand activity was ranked with predict_ligand_activities by corrected AUPR for both sender-agnostic and sender-filtered rankings, and regulatory potentials were taken from the ligand-target matrix. For visualization, ligand–TF regulatory potential scores below the 33$^{rd}$ percentile of the selected ligand-target link-weight distribution were set to zero using “prepare_ligand_target_visualization” with “cutoff = 0.33”; consequently, only candidate TFs with links passing this visualization threshold are shown in the heatmap, although all 33 candidate TFs were included in the NicheNet analysis. Absence from the heatmap does not imply zero regulatory potential. Dkk3 ranked highly and showed predicted regulatory potential toward Mitf; this was treated as exploratory rather than causal. These analyses supported Supplementary Figure 5.

#### Extended characterization of the rHSC transcriptional state

Apoptosis, core senescence, and SASP-related gene sets were evaluated in HSC_1 cells and rHSCs. Per-cell scores were the mean log-normalized expression of genes present in each set, without additional normalization or background-gene correction. The apoptosis set comprised Bax, Bbc3, Pmaip1, Bcl2l11, Casp3, Casp8, and Fas; the senescence-core set comprised Cdkn1a, Cdkn2a, Cdkn2b, Glb1, Serpine1, and Gdf15; and the SASP-related set comprised Il6, Cxcl1, Cxcl2, and Mmp3. Marker expression and scores were compared descriptively.

To assess transcriptional remodeling of rHSCs across fibrosis regression, rHSCs were stratified by experimental stage and per-cell scores were calculated for apoptosis-related, core senescence, activated HSC (aHSC), and quiescent HSC (qHSC)-associated gene sets using the same mean log-normalized expression approach. The aHSC-associated set comprised Col1a1, Acta2, Timp1, Loxl2, Tgfbr1, Pdgfrb, Vim, and Spp1. The qHSC-associated set comprised Gfap, Plin2, Adipor1, Pparg, Insig1, Foxo1, Cebpd, Ngfr, and Bambi. Stage-wise distributions were examined descriptively in fibrosis, recovery week 1, and recovery week 4 rHSCs. No cell-level hypothesis testing was used for these comparisons.

For rHSC GO analysis, differential expression between rHSCs and HSC_1 cells was assessed using Seurat FindMarkers with the default Wilcoxon rank-sum test, min.pct = 0.1, logfc.threshold = 0.25, and only.pos = FALSE. Genes with Bonferroni-adjusted P <0.05 and positive log fold change were analyzed with clusterProfiler::enrichGO using all genes in the HSC object as the background, org.Mm.eg.db, keyType = SYMBOL, pAdjustMethod = BH (Benjamini–Hochberg correction), pvalueCutoff = 0.05, and qvalueCutoff = 0.20. Terms were ranked by adjusted P value. These analyses supported Supplementary Figure 3.

**Extended processing and reference mapping of GSE295293**

Raw matrices from the nine GSE295293 samples were processed independently in Seurat. CreateSeuratObject used min.cells = 3 and min.features = 200. Cells were retained with >200 and <4,000 detected genes, <20,000 RNA counts, and <20% mitochondrial transcripts. Each sample was log-normalized; 3,000 variable genes were selected, followed by scaling and PCA.

Samples were integrated by reciprocal PCA using 3,000 integration features and the first 30 principal components for anchor identification and IntegrateData. The integrated object was scaled and analyzed by PCA, nearest-neighbor graph construction, UMAP, and clustering with 30 principal components and resolution 0.6. Major cell types were annotated with established markers. Extracted HSCs were reanalyzed in the RNA assay after log normalization and selection of 3,000 variable genes; PCA used 30 components, nearest-neighbor graph and UMAP used the first 20 components, and clustering used resolution 0.23.

For reference mapping, the query was log-normalized and 3,000 variable features were selected. FindTransferAnchors used the RNA assay, saved primary-study PCA reference, and dimensions 1-30. HSC cluster identity and binary rHSC/other-HSC labels were transferred with TransferData and MapQuery, and cells were projected onto the saved reference UMAP. The independent top-30 signature was scored with AddModuleScore using nbin = 24, ctrl = 100, and seed = 260703. Wt1 and Mdk detection was defined as normalized expression >0. These analyses supported Supplementary Figure 6.

#### Cross-species evaluation in human liver datasets

GSE136103 and GSE202379 were used to evaluate the murine rHSC-associated program in human liver mesenchymal cells. The CORE signature comprised 7 genes (Wt1, Mdk, Mitf, Hgf, Igf1, Vegfc, and Ngf), and the FULL signature comprised 882 genes significantly upregulated in murine rHSCs relative to HSC_1 cells and was defined without optimization against the human data. Mouse-to-human ortholog mapping was performed on September 10, 2026, using biomaRt v2.68.0 and the Ensembl BioMart mmusculus_gene_ensembl dataset at the live Asia endpoint (https://asia.ensembl.org:443/biomart/martservice?redirect=no). Mouse symbols were queried with external_gene_name, and human symbols were retrieved from hsapiens_homolog_associated_gene_name. Only ortholog_one2one mappings with hsapiens_homolog_orthology_confidence = 1 were accepted. One-to-many, many-to-many, low-confidence one-to-one, and unresolved mappings were excluded; duplicate accepted human symbols arising from different mouse genes were also excluded if present. No manual gene-symbol substitution, capitalization-based conversion, HGNC, HomoloGene, g:Profiler, orthogene, or pre-existing ortholog table was used for the original conversion. Because the live endpoint was queried, the exact Ensembl release was not recorded.

The mapping yielded 7 accepted high-confidence human orthologs for the 7-gene CORE signature and 805 for the 882-gene FULL signature. In GSE136103, the BioMart mapping was performed directly and genes absent from the scoring feature space were excluded, leaving 7 CORE and 776 FULL genes. For GSE202379, the locked FULL human ortholog mapping generated for GSE136103 was reused without rerunning conversion, and CORE was reused as the fixed human symbol list WT1, MDK, MITF, HGF, IGF1, VEGFC, and NGF. Genes absent from the SCT feature space were omitted, leaving 7 CORE and 747 FULL genes. Final Supplementary Figure 7 assembly reused finalized CSV outputs and locked signatures and did not rerun ortholog conversion, module scoring, mouse differential-expression gene selection, or statistical testing.

Seurat AddModuleScore used nbin = 24, ctrl = 100, and seed = 20260910 for core and broad signatures. Analyses focused on hepatic mesenchymal and stellate populations and used donor-level comparisons when donor information was available. In GSE136103, scores were evaluated in HSCs, vascular smooth muscle cells, scar-associated mesenchymal cells, and mesothelial cells. GSE202379 contained 47 donors in total. Primary donor-level analyses were restricted to the 42 donors with at least 10 annotated stellate cells and compared the donor-level clinical groups. Overall differences were assessed using the Kruskal–Wallis test separately for CORE and FULL; when the overall P value was <0.05, all pairwise clinical-group comparisons were tested. Pairwise post-hoc comparisons used two-sided Wilcoxon rank-sum tests (wilcox.test, exact = FALSE, with the default continuity correction), with Benjamini–Hochberg adjustment across the 10 pairwise comparisons among the five clinical groups separately for CORE and FULL. Individual core genes were also examined; because WT1 and MDK were low or context-dependent, cross-species correspondence was interpreted at the broader program level. These analyses supported Supplementary Figure 7.

**Pharmacological inhibition of MDK**

Male mice received TAA (300 mg/L) in drinking water for 8 weeks, followed by TAA withdrawal and daily administration of the midkine inhibitor iMDK for 2 weeks. iMDK (Merck Millipore; catalog no. 5.08052.0001; lot 3808865) was prepared as a 5 mg/mL stock by dissolving 10 mg iMDK in 2 mL DMSO. The stock was divided into 50-µL aliquots and stored at −30°C. Immediately before administration, 50 µL stock was mixed with 731.25 µL corn oil. Vehicle-treated control mice received the corresponding DMSO/corn oil vehicle without iMDK. Each group contained six mice (n = 6 per group).

iMDK was administered intraperitoneally once daily at approximately 10:00 AM for 2 weeks (14 administrations) at a total weekly dose of 9 mg/kg body weight, corresponding to approximately 1.29 mg/kg/day. The dose was based on Masui et al., 2016 (1). Isoflurane anesthesia was used for procedures requiring anesthesia. P values for two-group comparisons in the iMDK experiment were calculated using two-sided unpaired Student’s t-tests.

##### Single-factor MDK overexpression

After 12 weeks of TAA treatment, male mice received rAAV-GFP or rAAV-artMdk at $1 \times 10^{11}$ vector genomes per mouse by intravenous injection and were analyzed 2 weeks after vector administration. Five mice per group were included in the final quantitative analyses. One rAAV-artMdk-treated mouse was excluded because vector-derived artMdk expression was not detected. Hepatic transgene expression was assessed by quantitative PCR using primers specific to the synthetic artMdk sequence, allowing vector-derived artMdk expression to be distinguished from endogenous Mdk. Isoflurane anesthesia was used for procedures requiring anesthesia. Relative gene-expression values from individual mice in Supplementary Figure 8I were visualized in R using ggplot2. Violin plots were generated with `geom_violin(trim = FALSE)`; individual animals were overlaid as jittered points, and group means were indicated by white diamonds. P values for qPCR measurements in Supplementary Figure 8I were calculated using two-sided Student’s t-tests.

##### Western blotting

Whole-liver proteins were extracted in TNE buffer containing 20 mM Tris-HCl (pH 7.4), 150 mM NaCl, 2 mM EDTA, 1% Nonidet P-40, and Protease Inhibitor Cocktail Set III

(Calbiochem; cat. no. 539134; lot D00125001) added at 1:1,000. Protein (20 µg in 20 µL per lane) was separated on a 20% polyacrylamide gel at 65 V for 20 min and then 100 V for 90 min. Proteins were transferred to an Immobilon-P PVDF membrane (Millipore; IPVH00010) by tank transfer at 100 mA for 60 min, followed by Ponceau S staining.

Membranes were blocked with 2% skim milk in PBS-T for 30 min at room temperature and incubated overnight at 4°C with goat anti-actin antibody (Actin [I-19], Santa Cruz Biotechnology, sc-1616, lot G1615; 1:1,000) or sheep anti-mouse MDK antibody (R&D Systems, AF7769; 1:25). The corresponding HRP-conjugated secondary antibodies were Polyclonal Rabbit Anti-Goat Immunoglobulins/HRP (Ref. P0160, lot 00039881; 1:5,000) and rabbit anti-sheep IgG (H+L) secondary antibody, HRP (Zymed/Thermo Fisher Scientific, 61-8620; 1:5,000). Secondary antibodies were incubated for 1 h at room temperature. Signals were detected with Immobilon Forte Western HRP Substrate (WBLUF0100) and imaged using Fusion FX (Vilber Bio Imaging).

**GFP and MDK visualization**

Liver tissue was fixed in Mildform 10N (FUJIFILM Wako; 133-10311), paraffin-embedded, and sectioned at 6 µm. For MDK staining, sections underwent proteinase K retrieval (1 mg/mL in PBS, 37°C, 15 min), Block Ace treatment for 2 h, incubation with sheep anti-mouse MDK antibody (R&D Systems, AF7769; 1:100) overnight, and incubation with donkey anti-sheep IgG NL557 (R&D Systems, NL010; 1:500) for 2 h. Nuclei were stained with Hoechst 33342. GFP was detected by endogenous fluorescence. These stains were used for representative visualization and were not quantified.

1. Masui M, Okui T, Shimo T, Takabatake K, Fukazawa T, Matsumoto K, et al. Novel Midkine Inhibitor iMDK Inhibits Tumor Growth and Angiogenesis in Oral Squamous Cell Carcinoma. Anticancer Res. 2016;36:2775–2781.

### Supplementary Table 1. Antibodies and detection reagents used in this study.

| Target or reagent | Host or clone | Conjugate | Supplier | Catalog no. | Lot no. | Application | Dilution | Note |
|---|---|---|---|---|---|---|---|---|
| PECAM1 (CD31) | Rat, MEC13.3 | PE | BD Biosciences | 553373 | 49700 | FACS | 1:100 | LSEC identification |
| F4/80 | Rat, BM8 | Biotin | BioLegend | 123106 | B157667 | FACS | 1:100 | Macrophage identification |
| Streptavidin | — | APC | BD Pharmingen | 554067 | 7040924 | FACS | 1:300 | Detection of biotinylated F4/80 antibody |
| EpCAM | In-house antibody | FITC | In-house | — | — | FACS | 1:100 | Prepared and FITC-conjugated in-house; antibody described in Okabe et al., Development (2009), doi:10.1242/dev.031369 |
| MDK | Sheep anti-mouse | — | R&D Systems | AF7769 | CHHP0122081 | Immunohistochemistry | 1:100 | Primary antibody |
| Sheep IgG | Donkey anti-sheep | NL557 | R&D Systems | NL010 | — | Immunohistochemistry | 1:500 | Secondary antibody for MDK detection |
| MDK | Sheep anti-mouse | — | R&D Systems | AF7769 | CHHP0122081 | Western blotting | 1:25 | Primary antibody |
| Sheep IgG (H+L) | Rabbit anti-sheep | HRP | Zymed / Thermo Fisher Scientific | 61-8620 | 30678141R | Western blotting | 1:5,000 | Secondary antibody for MDK detection |
| Actin | Goat, I-19 | — | Santa Cruz Biotechnology | sc-1616 | G1615 | Western blotting | 1:1,000 | Loading control |
| Goat immunoglobulins | Rabbit anti-goat, polyclonal | HRP | Dako | P0160 | 00039881 | Western blotting | 1:5,000 | Secondary antibody for actin detection |

#### Supplementary Table 2. Primer sequences used for quantitative real-time PCR

| Target | Forward primer (5′–3′) | Reverse primer (5′–3′) |
|---|---|---|
| eGFP | AGTCCGCCCTGAGCAAAGA | TCCAGCAGGACCATGTGATC |
| Igf1 | TGACCGCACCTGCAATAAAGA | GTCCAGCTGTGGTGGAGCTG |
| artMdk | GGCATGGGTTTTCGTGAAGG | AGCTTTTGTGCCTGTTGAGC |
| Vegfc | CTTCGAGTCGGGACTGGGCT | TTTGCCTTCAAAAGCCTTGACCT |
| Ngf | GGGAGCGCATCGAGTGACTT | TCACTGGCTTGAGGCACAGC |
| Hgf | GGAGCCACAAGGATCTGGAGC | GCCGGGCTGAAAGAATCAAAGC |
| Col1a1 | TCGTGGCTTCTCTGGTCTC | CCGTCTTTGCCAGGAGAA |
| Col3a1 | TCCCCTGGAATCTGTGAATC | TGAGTCGAATTGGGGAGAAT |
| Tgfb1 | TCAGACATTCGGGAAGCAGT | ACGCCAGGAATTGTTGCTAT |
| Timp1 | GCAAAGAGCTTTCTCAAAGACC | AGGGATAGATAAACAGGGAAACACT |
| Mmp9 | TGACTACGATAAGGACGGCAAA | AGATGAACGGGAACACACAGG |
| Il6 | GGACTTAGCTCGTCTCATTCATTC | TGTAAATCTTTTACCTAAAGGAGGAAA |
| Acta2 | GACACCACCCACCCAGAGT | ACATAGCTGGAGCAGCGTCT |
| Actb | CCAACCGTGAAAAGATGACC | CCAGAGGCATACAGGGACAG |

*artMdk primers specifically amplify the synthetic vector-derived artMdk sequence and do not detect endogenous Mdk.*

#### SUPPLEMENTARY FIGURE LEGENDS

**Supplementary Figure 1. Hallmark pathway changes during spontaneous fibrosis regression.**

A, Top Hallmark gene sets enriched in fibrotic liver (TAA8w) relative to recovery 4 weeks, with normalized enrichment score (NES). Negative NES values indicate enrichment in TAA8w.

B, Top Hallmark gene sets enriched at recovery 4 weeks relative to TAA8w. Positive NES values indicate enrichment during recovery.

C, Dot plot of Hallmark pathways enriched in TAA8w. Dot size indicates the number of genes contributing to the gene set, and color indicates the adjusted P value. Pathways associated with cell-cycle progression and proliferative signaling, including G2M checkpoint, E2F targets, MYC targets, and mTORC1 signaling, were preferentially enriched in fibrotic liver.

D, Dot plot of Hallmark pathways enriched at recovery 4 weeks. Dot size indicates the number of genes contributing to the gene set, and color indicates the adjusted P value. Recovery was associated with enrichment of pathways related to coagulation, bile acid metabolism, xenobiotic metabolism, peroxisomal function, and complement.

**Supplementary Figure 2. Cell-type-specific transcriptional remodeling during fibrosis regression.**

A, Differential gene-expression analysis of endothelial cells (ECs) between healthy liver and recovery 1 week. Recovery-associated genes included scavenger/endocytic receptors such as Stab2, Lyve1, and Cd36. Gene Ontology enrichment analysis of recovery-upregulated genes highlighted processes including receptor-mediated endocytosis, cell migration, and cell–substrate adhesion.

B, Differential expression between Kupffer cells (KC) and monocyte-derived macrophages (MDM) at recovery week 4. Candidate MDM-enriched surface-associated genes, including Alcam, Sema4d, and Slamf7, are highlighted. Density plots show the distribution of selected macrophage-associated genes in KC and MDM pooled across healthy, fibrotic, recovery 1-week, and recovery 4-week samples. These candidates were identified transcriptionally and were not functionally validated as isolation markers.

C, Differential gene-expression analysis of cholangiocytes between healthy liver and recovery 4 weeks. Recovery-associated genes included multiple apoptosis-related genes, and Gene Ontology analysis highlighted regulation of apoptotic signaling together with RNA-regulatory processes.

**Supplementary Figure 3. Detailed transcriptional characterization of the rHSC state.**

A, Proportion of rHSCs (HSC_2) among HSC_1 and HSC_2 cells in healthy liver, fibrosis, and recovery weeks 1 and 4.

B, Dot plot comparing quiescent HSC-associated, activated/fibrotic HSC-associated, rHSC-associated, apoptosis-related, core senescence, and senescence-associated secretory phenotype (SASP)-related genes between rHSCs and HSC_1 cells. Dot size indicates the percentage of cells expressing each gene, and color indicates scaled average expression.

C, Density-based visualization of Wt1 and Mdk expression across the HSC UMAP, showing preferential localization of these genes within the rHSC compartment.

D, Mean log-normalized expression scores for apoptosis-related, core senescence, and SASP-related gene sets in rHSCs and HSC_1 cells, assessing whether rHSCs were associated with a generalized apoptotic or senescence-like transcriptional program.

E, Stage-dependent changes in transcriptional programs within rHSCs. Violin plots show per-cell mean log-normalized expression scores for apoptosis-related, core senescence, activated HSC (aHSC), and quiescent HSC (qHSC)-associated gene sets in rHSCs during fibrosis and at recovery weeks 1 and 4. Apoptosis-related and aHSC scores decreased during recovery, whereas the qHSC-associated score was modestly higher at recovery week 4 than during fibrosis. The core senescence score decreased at recovery week 1 and partially increased at recovery week 4. These patterns indicate transcriptional remodeling of rHSCs during fibrosis regression rather than simple persistence of the fibrotic activated state. Boxes indicate the median and interquartile range.

F, Gene Ontology Biological Process enrichment analysis of genes upregulated in rHSCs relative to HSC_1 cells. Enriched terms included extracellular matrix organization, connective tissue development, TGF-β-related signaling, cell–substrate adhesion, and developmental processes. Dot size indicates gene count and color indicates adjusted P value.

**Supplementary Figure 4. Inferred HSC-centered intercellular signaling persists during late fibrosis regression.**

A, CellChat-inferred intercellular communication network at recovery 4 weeks among cholangiocytes (CHOL), HSC_1, rHSCs, Kupffer cells (KC), liver sinusoidal endothelial cells (LSEC), and monocyte-derived macrophages (MDM).

B, Outgoing and incoming interaction strengths of the displayed major non-parenchymal cell populations at recovery 4 weeks. Cholangiocytes were omitted from this scatterplot because their persistently high SPP1-dominated inferred outgoing signal obscured comparison among the other populations. rHSCs remained a prominent source of inferred outgoing signaling among the populations shown during late regression.

C, Differential CellChat analysis comparing recovery 4 weeks with healthy liver. Signaling pathways are positioned according to changes in inferred outgoing and incoming interaction strength, highlighting persistent recovery-associated inferred signaling pathways including MK, VEGF, TGF-β, BMP, GAS, and SPP1 signaling. MK is the CellChat pathway label for midkine (MDK) signaling.

D, CellChat-inferred MK signaling network at recovery 4 weeks, showing predicted cellular sources and targets of MK-mediated communication.

E, DEG-mapped inferred ligand–receptor interactions from rHSCs to cholangiocytes, HSC_1, KC, LSEC, and MDM associated with recovery week 4. Dot color represents inferred communication probability, and interactions were retained according to the specified differential-expression and ligand fold-change criteria. Representative interactions include Hgf–Met, Igf1–Igf1r, Vegfc–Vegfr family receptors, Mdk receptor interactions, and GAS-, BMP-, TGF-β-, and galectin-associated signaling.

F, Quantitative summary of the top rHSC-derived ligands showing increased inferred signaling at recovery 4 weeks relative to healthy liver. Ligands were ranked by mean ligand log fold change among F4-upregulated rHSC-derived CellChat interactions. Components of the rHSC-associated factor mixture meeting the predefined F4-upregulation criterion are highlighted, including Hgf, Igf1, Mdk, and Vegfc; Ngf did not meet this cutoff at recovery 4 weeks.

G, Number of inferred CellChat interactions between HSCs and LSECs across healthy (H), fibrotic (Fib), recovery 1-week (R1), and recovery 4-week (R4) conditions, showing stage-dependent changes in inferred bidirectional communication between these populations during fibrosis and regression.

H, CellChat-inferred signaling interactions from HSCs to LSECs and HSCs across healthy (He), fibrotic (Fib), recovery 1-week (R1w), and recovery 4-week (R4w) conditions. Selected pathways include VEGF, MK, FGF, WNT, and BMP signaling. Dot color indicates inferred communication probability. Only inferred ligand–receptor interactions with CellChat P < 0.05 at the communication-probability inference step are shown.

I, CellChat-inferred signaling interactions from LSECs to HSCs and LSECs across the four experimental conditions. Selected pathways include GALECTIN, VISFATIN, GAS, and BMP signaling, showing predicted cross-population and within-lineage signaling. Only inferred ligand–receptor interactions with CellChat P < 0.05 at the communication-probability inference step are shown.

**Supplementary Figure 5. Downstream refinement and upstream ligand inference for Wt1/Mitf-associated rHSC programs.**

A–C, Refinement of CellOracle-predicted Wt1/Mitf targets using HSC gene co-expression information. A, Schematic of the target-refinement strategy. CellOracle-predicted Wt1 and Mitf targets retained after filtering at P < 0.001 and selecting the top 2,000 edges by absolute coefficient per condition were intersected with genes in co-expression modules containing both Wt1 and Mitf. Four genes, Gas7, Plxna4, Runx1, and Serpine2, were shared by both transcription factors at both recovery stages. B, Violin plots showing the expression of the four shared candidate target genes across healthy, fibrotic, recovery 1-week, and recovery 4-week HSCs. C, CellOracle-predicted candidate regulatory relationships between Wt1/Mitf and the four shared candidate targets at recovery 1 week. All eight transcription factor (TF)-to-target edges passed the filters used in A; the additional Wt1-to-Mitf edge is shown from the raw GRN. Edge color indicates the source transcription factor, and edge width represents the absolute value of the CellOracle coef_mean. All displayed coefficients were positive. These inferred relationships and co-expression-based refinements do not establish direct transcriptional regulation.

D–F, NicheNet analysis to nominate candidate upstream ligands associated with the recovery-related HSC transcription factor program. (35) D, Ligand activity scores (corrected area under the precision–recall curve (AUPR)) for candidate upstream ligands predicted to regulate the selected HSC transcription factor program. E, Expression of candidate ligands across the analyzed sender cell populations. F, For visualization, regulatory-potential scores below the 33rd percentile of selected ligand-target link-weight distribution were set to zero.

**Supplementary Figure 6. External validation of an rHSC-like transcriptional state using an independent liver fibrosis regression dataset.**

A, UMAP of the GSE295293 liver single-cell dataset, colored by major cell type annotation.

B, Feature plots showing representative lineage markers used to identify major hepatic cell populations: Alb, Epcam, Kdr, Adgre1, and Lrat.

C, UMAP of the HSC subset after reclustering, colored by HSC subcluster.

D, Distribution of the HSC subcluster across control (CONT), TAA-induced fibrosis (TAA), and regression (TAA_RE) conditions.

E, Projection of GSE295293 HSCs onto the primary HSC reference UMAP, with all conditions combined. Query cells classified by reference mapping as rHSC-like (n = 1,167) or other HSCs (n = 3,423) are highlighted in red and blue, respectively. Reference cells are shown in gray.

F, Independent rHSC signature scores in GSE295293 HSCs, classified as rHSC-like or other HSCs, with all conditions combined. Mapped rHSC-like cells showed higher signature scores than other HSCs. Boxes indicate the median and interquartile range.

G, Dot plot showing Wt1 and Mdk expression in mapped rHSC-like and other HSCs from GSE295293, with all conditions combined. Dot size indicates the percentage of cells expressing each gene, and color indicates mean expression. Both genes were enriched in the mapped rHSC-like population.

H, Numbers and proportions of GSE295293 HSCs mapped to an rHSC-like state using the primary HSC reference for each sample.

I, Proportion of cells mapped to an rHSC-like state among GSE295293 HSCs in each condition.

J, Summary of reference-mapping inputs and transfer anchors, including shared reference variable features, common RNA genes, and the numbers of query cells, reference cells, and transfer anchors.

**Supplementary Figure 7. Cross-species evaluation of the murine rHSC-associated transcriptional program in human liver datasets.**

(A) Overview of the evaluation of the locked murine CORE signature (WT1, MDK, MITF, HGF, IGF1, VEGFC and NGF; human symbols shown) and FULL rHSC-up program after mouse-to-human ortholog conversion in two independent human liver datasets. Previously completed scores and original author-defined populations were retained without signature optimization or reannotation.

(B) Previously computed CORE and FULL module scores in GSE136103 author-defined vascular smooth muscle cells (VSMC), hepatic stellate cells (HSC), scar-associated mesenchymal cells (SAMes) and Mesothelia. Each point represents the mean score for one donor-population combination; colors distinguish healthy and cirrhotic donors. All available combinations were retained, with 8, 9, 10 and 4 donors represented in VSMC, HSC, SAMes and Mesothelia, respectively; no minimum cell-count threshold was imposed. The median of donor-mean scores is highest in SAMes for both signatures. These population labels do not establish equivalence to murine rHSC.

(C,D) Previously computed donor-mean CORE (C) and FULL (D) scores in GSE202379 author-defined Stellate cells. One point represents one donor. GSE202379 comprised 47 donors in total; primary donor-level analyses in panels C and D were restricted to the 42 donors with at least 10 annotated Stellate cells: healthy control, n = 4; Non-alcoholic Fatty Liver Disease (NAFLD), n = 6; Non-alcoholic Steatohepatitis (NASH) without cirrhosis, n = 23; NASH with cirrhosis, n = 4; and end stage, n = 5. Overall Kruskal-Wallis P values are 0.0300 and 0.0233, respectively, taken from the completed analysis. Pairwise post-hoc comparisons used two-sided Wilcoxon rank-sum tests (wilcox.test, exact = FALSE, with the default continuity correction), with Benjamini–Hochberg adjustment across the 10 pairwise comparisons among the five clinical groups separately for CORE and FULL. No Benjamini-Hochberg-adjusted pairwise comparison was significant. Stage groups are cross-sectional, and the non-monotonic patterns are not a longitudinal trajectory or a simple measure of fibrosis severity.

In B, C and D, boxes indicate the median and interquartile range, with whiskers extending to the most extreme observations within 1.5 times the interquartile range; all donor points, including outliers, are shown. Scores, population definitions and statistical results were reused from the finalized analyses; no new biological analysis was performed for figure assembly. Together, the completed analyses support partial, context-dependent cross-species conservation of an rHSC-associated transcriptional program, without establishing a directly equivalent human rHSC population, a conserved functional role, or conservation of WT1-MDK expression in human Stellate cells.

**Supplementary Figure 8. Effects of pharmacological MDK inhibition and hepatic MDK overexpression during regression of TAA-induced liver fibrosis.**

A–C, Male mice were treated with TAA for 8 weeks, followed by once-daily intraperitoneal administration of iMDK for 2 weeks (14 administrations; 9 mg/kg/week). Vehicle-treated controls received Dimethyl sulfoxide (DMSO)/corn oil. DMSO, n = 6; iMDK, n = 6.

A, Representative Sirius Red staining. Scale bar, 200 μm.

B, Quantification of Sirius Red-positive area. Bars show mean ± Standard Error of the Mean (SEM). The P value was calculated using a two-sided Student's t-test.

C, Hepatic expression of Acta2, Tgfb1, Col1a1, Col3a1, and Timp1 measured by qPCR. Violin plots show individual-animal distributions; points represent individual mice and white diamonds indicate group means.

D,E, and G–I, Male mice were treated with TAA for 12 weeks, followed by intravenous administration of rAAV-GFP or rAAV-artMdk (1 × 10^11 vector genomes/mouse) and analyzed 2 weeks later. Quantitative analysis in D, H, and I included five mice per group; one rAAV-artMdk-treated mouse was excluded because vector-derived artMdk expression was not detected.

D, Hepatic expression of enhanced green fluorescent protein (eGFP) and vector-derived artMdk measured by qPCR. N.D., not detected. Statistical comparisons were not performed for transgene expression when expression was undetectable in one of the groups.

E, Western blot analysis of MDK and Actin. Animal IDs are shown above each lane. No. 80 is shown for transparency but was excluded from quantitative analyses because vector-derived artMdk expression was not detected. Cropped blots are shown; uncropped source blots were retained.

F, Representative GFP and MDK localization in liver sections from a separate cohort (n = 7 mice per group) subjected to the same TAA treatment, rAAV administration, and sampling schedule described above. GFP expression was assessed by endogenous fluorescence. Scale bar, 200 μm.

G, Representative Sirius Red staining from rAAV-GFP and rAAV-artMdk mice. Scale bar, 200 μm.

H, Quantification of Sirius Red-positive area. Bars show mean ± SEM. The P value was calculated using a two-sided Student's t-test.

I, Relative expression of Acta2, Col1a1, Col3a1, Tgfb1, and Timp1 measured by qPCR. Violin plots show individual-animal distributions; points represent individual mice and white diamonds indicate group means. P values for qPCR measurements in C and I were calculated using two-sided Student's t-tests.

**A**

| ID | NES | pvalue | p.adjust | size |
|---|---|---|---|---|
| G2M_checkpoint | -3.8326626 | 1.00E-10 | 1.13E-09 | 94 |
| E2F_targets | -3.4355869 | 1.00E-10 | 1.13E-09 | 96 |
| MTORC1_signaling | -3.1152132 | 1.00E-10 | 1.13E-09 | 78 |
| Myc_targets_v1 | -3.0953379 | 1.00E-10 | 1.13E-09 | 84 |
| Mitotic_spindle | -2.768466 | 1.36E-08 | 1.23E-07 | 75 |
| Cholesterol_homeostasis | -2.863019 | 2.11E-08 | 1.58E-07 | 47 |
| Myc_targets_v2 | -2.6252074 | 9.71E-07 | 5.46E-06 | 34 |
| Androgen_response | -2.3234666 | 0.00011224 | 0.0005612 | 36 |
| Estrogen_response_early | -1.959368 | 0.0013264 | 0.00497399 | 59 |
| Estrogen_response_late | -1.9311824 | 0.00152111 | 0.00526537 | 58 |

**B**

| ID | NES | pvalue | p.adjust | size |
|---|---|---|---|---|
| Coagulation | 2.58685782 | 5.52E-08 | 3.55E-07 | 57 |
| Bile_acid_metabolism | 1.91978835 | 0.00094425 | 0.00424911 | 51 |
| Xenobiotic_metabolism | 1.83895587 | 0.00115385 | 0.00472031 | 115 |
| Peroxisome | 1.62595816 | 0.0160769 | 0.03807686 | 38 |
| Complement | 1.55773559 | 0.01590737 | 0.03807686 | 62 |
| KRAS_signaling_dn | 1.71414546 | 0.01801345 | 0.03887062 | 28 |
| Interferon_gamma_response | 1.49367287 | 0.06140351 | 0.11052632 | 46 |
| Oxidative_phosphorylation | 1.39683156 | 0.08566434 | 0.14277389 | 49 |
| KRAS_signaling_up | 1.29081308 | 0.14642857 | 0.21964286 | 44 |
| Adipogenesis | 1.1299574 | 0.2722323 | 0.35515448 | 75 |

**C**

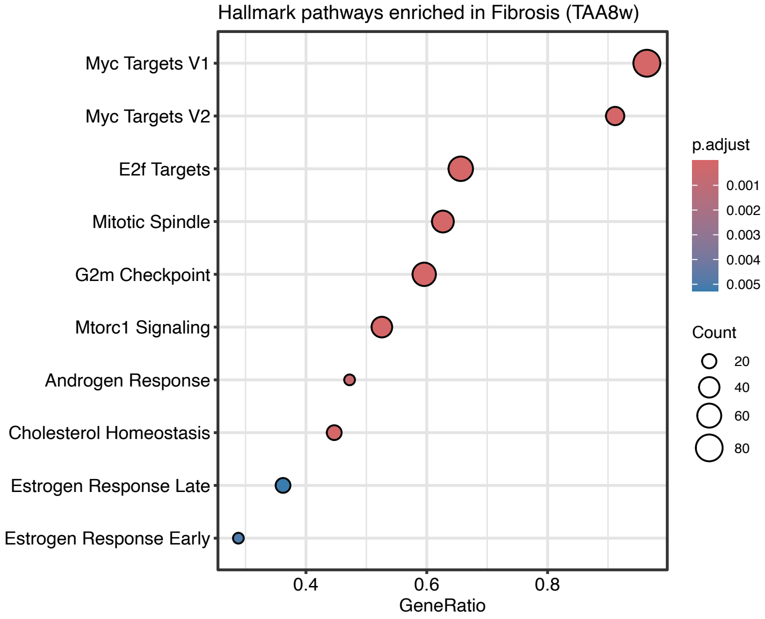


**D**

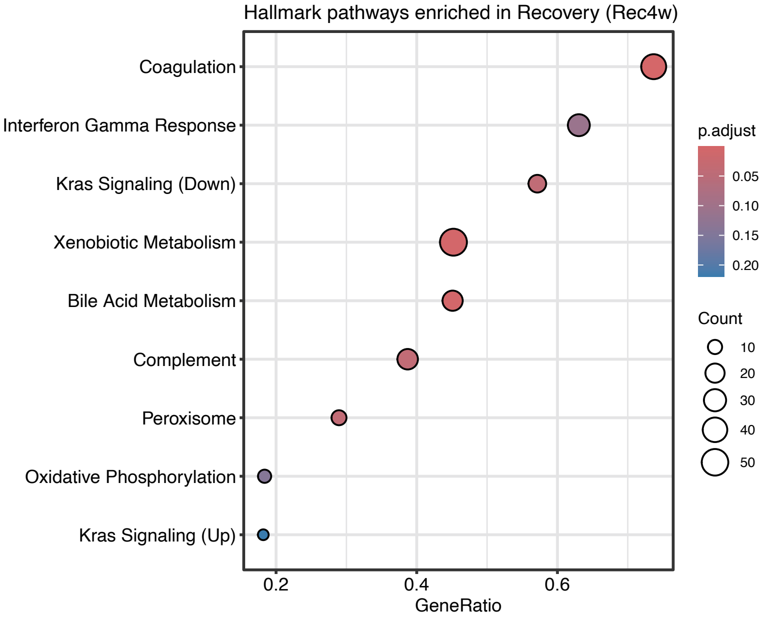


Transcriptomic pathway changes during spontaneous fibrosis regression.

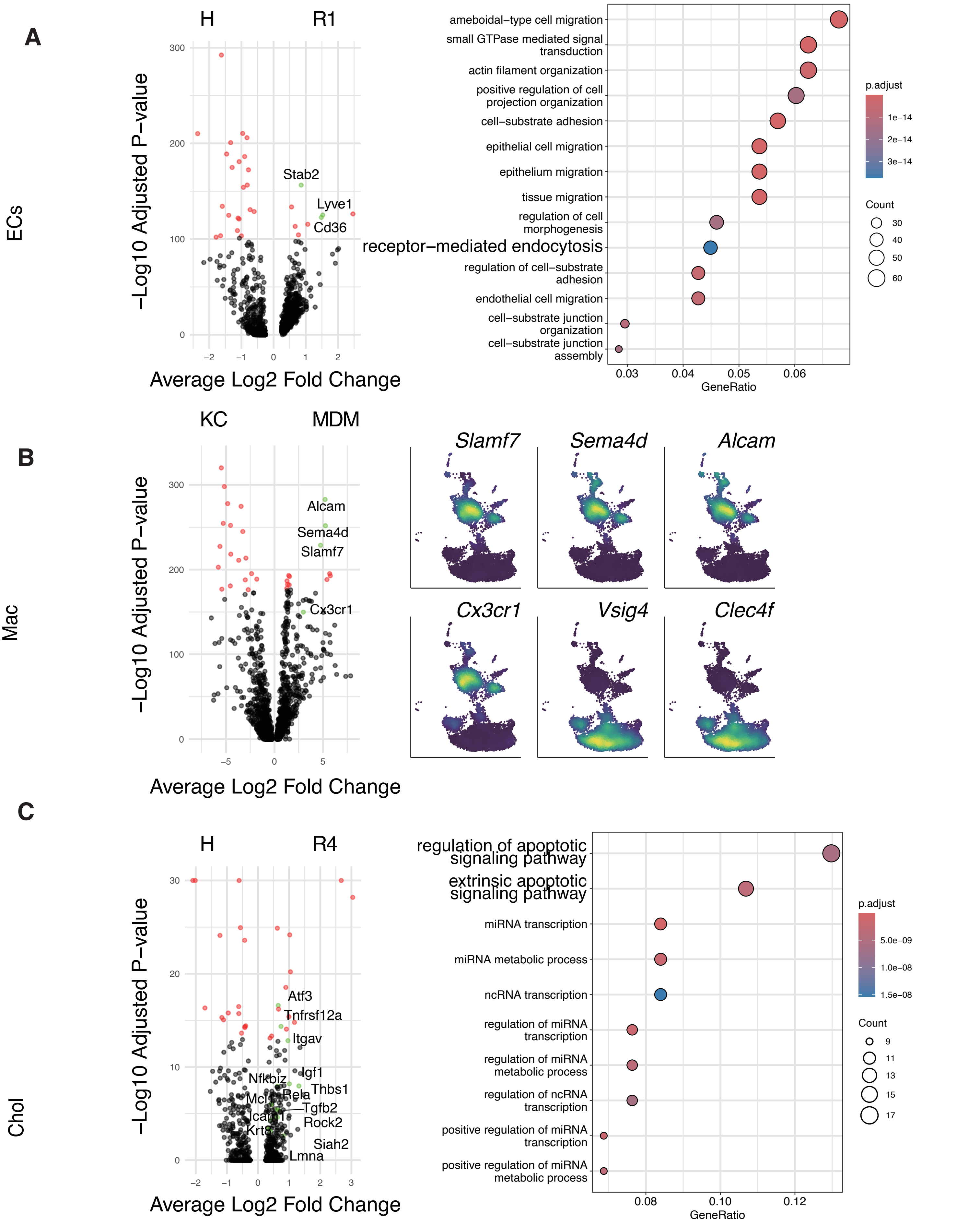


Cell-type-specific transcriptional remodeling during fibrosis regression.

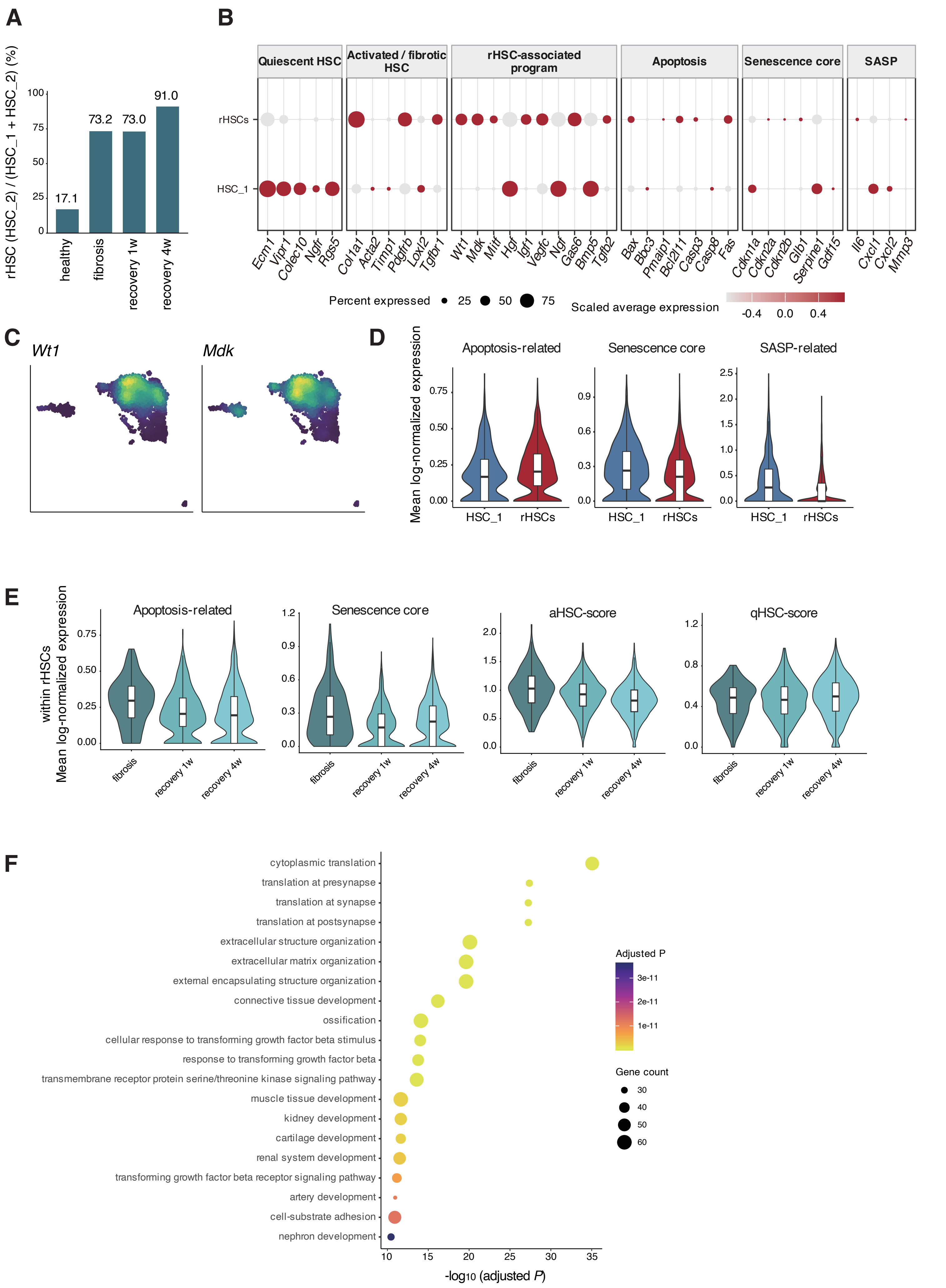
A
B
C
D
E
F
rHSC (HSC_2) / (HSC_1 + HSC_2) (%)
17.1
73.2
73.0
91.0
healthy
fibrosis
recovery 1w
recovery 4w
Quiescent HSC
Activated / fibrotic HSC
rHSC-associated program
Apoptosis
Senescence core
SASP
rHSCs
HSC_1
Ecm1 Vipr1 Colec10 Ngfr Rgs5
Col1a1 Acta2 Timp1 Pdgfrb Loxl2 Tgfbr1
Wt1 Mdk Mitf Hgf Igf1 Vegfc Ngf Gas6 Bmp5 Tgfb2
Bax Bbc3 Pmaip1 Bcl2l11 Casp3 Casp8 Fas
Cdkn1a Cdkn2a Cdkn2b Glb1 Serpine1 Gdf15
Il6 Cxcl1 Cxcl2 Mmp3
Percent expressed
Scaled average expression
Wt1
Mdk
Mean log-normalized expression
Apoptosis-related
Senescence core
SASP-related
within rHSCs
aHSC-score
qHSC-score
cytoplasmic translation
translation at presynapse
translation at synapse
translation at postsynapse
extracellular structure organization
extracellular matrix organization
external encapsulating structure organization
connective tissue development
ossification
cellular response to transforming growth factor beta stimulus
response to transforming growth factor beta
transmembrane receptor protein serine/threonine kinase signaling pathway
muscle tissue development
kidney development
cartilage development
renal system development
transforming growth factor beta receptor signaling pathway
artery development
cell-substrate adhesion
nephron development
Adjusted P
Gene count
-log10 (adjusted P)

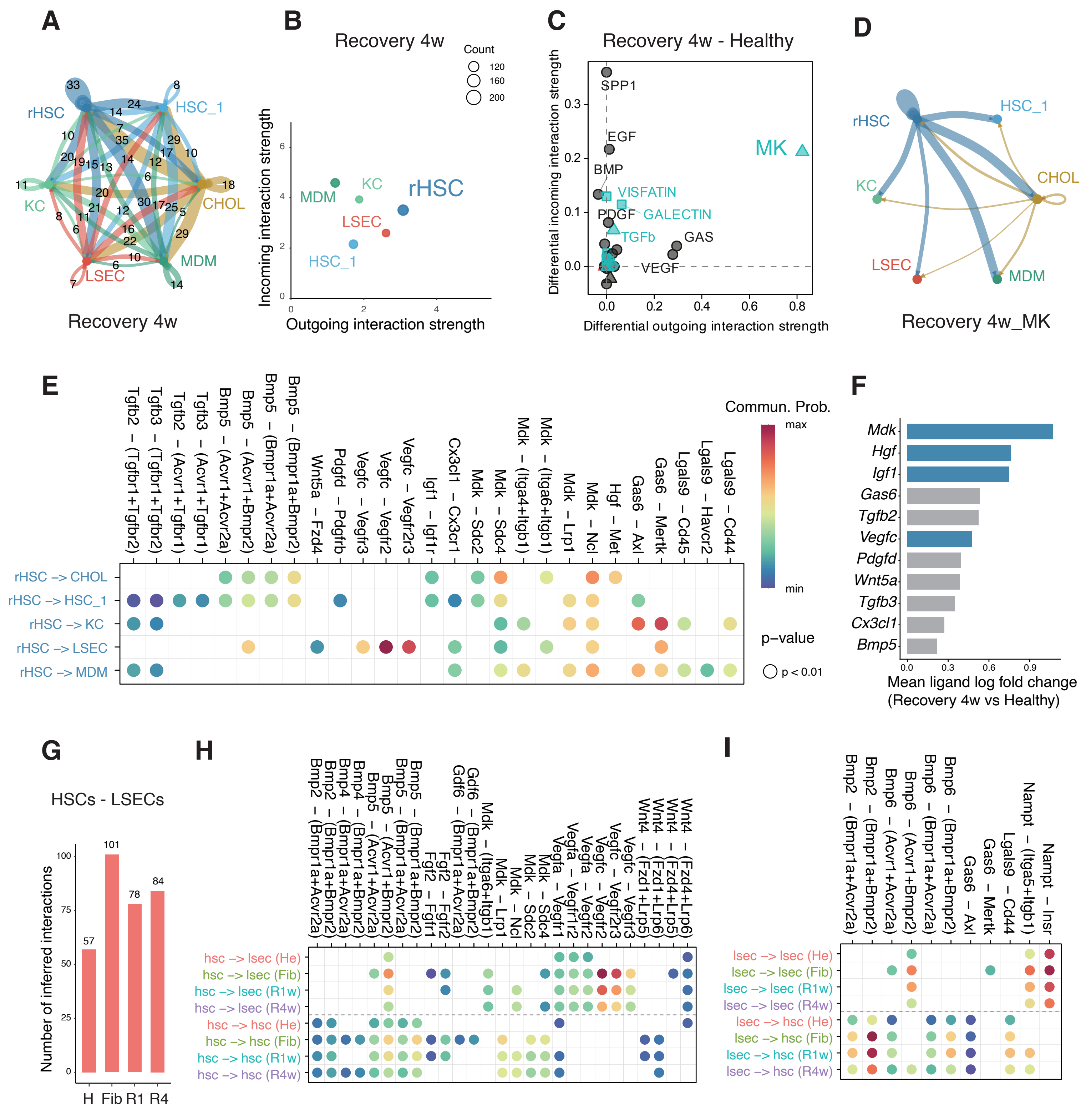
A
rHSC
HSC_1
KC
CHOL
LSEC
MDM
Recovery 4w
B
Recovery 4w
Count
120
160
200
Incoming interaction strength
Outgoing interaction strength
MDM
KC
rHSC
LSEC
HSC_1
C
Recovery 4w - Healthy
Differential incoming interaction strength
Differential outgoing interaction strength
SPP1
EGF
BMP
VISFATIN
PDGF
GALECTIN
TGFb
GAS
VEGF
MK
D
rHSC
HSC_1
KC
CHOL
LSEC
MDM
Recovery 4w_MK
E
Commun. Prob.
max
min
p–value
p < 0.01
rHSC –> CHOL
rHSC –> HSC_1
rHSC –> KC
rHSC –> LSEC
rHSC –> MDM
Tgfb2 – (Tgfbr1+Tgfbr2)
Tgfb3 – (Tgfbr1+Tgfbr2)
Tgfb2 – (Acvr1+Tgfbr1)
Tgfb3 – (Acvr1+Tgfbr1)
Bmp5 – (Acvr1+Acvr2a)
Bmp5 – (Acvr1+Bmpr2)
Bmp5 – (Bmpr1a+Acvr2a)
Bmp5 – (Bmpr1a+Bmpr2)
Wnt5a – Fzd4
Pdgfd – Pdgfrb
Vegfc – Vegfr3
Vegfc – Vegfr2
Vegfc – Vegfr2r3
Igf1 – Igf1r
Cx3cl1 – Cx3cr1
Mdk – Sdc2
Mdk – Sdc4
Mdk – (Itga4+Itgb1)
Mdk – (Itga6+Itgb1)
Mdk – Lrp1
Mdk – Ncl
Hgf – Met
Gas6 – Axl
Gas6 – Mertk
Lgals9 – Cd45
Lgals9 – Havcr2
Lgals9 – Cd44
F
Mdk
Hgf
Igf1
Gas6
Tgfb2
Vegfc
Pdgfd
Wnt5a
Tgfb3
Cx3cl1
Bmp5
Mean ligand log fold change
(Recovery 4w vs Healthy)
G
HSCs - LSECs
Number of inferred interactions
57
101
78
84
H
Fib
R1
R4
H
hsc –> lsec (He)
hsc –> lsec (Fib)
hsc –> lsec (R1w)
hsc –> lsec (R4w)
hsc –> hsc (He)
hsc –> hsc (Fib)
hsc –> hsc (R1w)
hsc –> hsc (R4w)
I
lsec –> lsec (He)
lsec –> lsec (Fib)
lsec –> lsec (R1w)
lsec –> lsec (R4w)
lsec –> hsc (He)
lsec –> hsc (Fib)
lsec –> hsc (R1w)
lsec –> hsc (R4w)
Bmp2 – (Bmpr1a+Acvr2a)
Bmp2 – (Bmpr1a+Bmpr2)
Bmp6 – (Acvr1+Acvr2a)
Bmp6 – (Acvr1+Bmpr2)
Bmp6 – (Bmpr1a+Acvr2a)
Bmp6 – (Bmpr1a+Bmpr2)
Gas6 – Axl
Gas6 – Mertk
Lgals9 – Cd44
Nampt – (Itga5+Itgb1)
Nampt – Insr

Saijou et al. Sup Figure 5

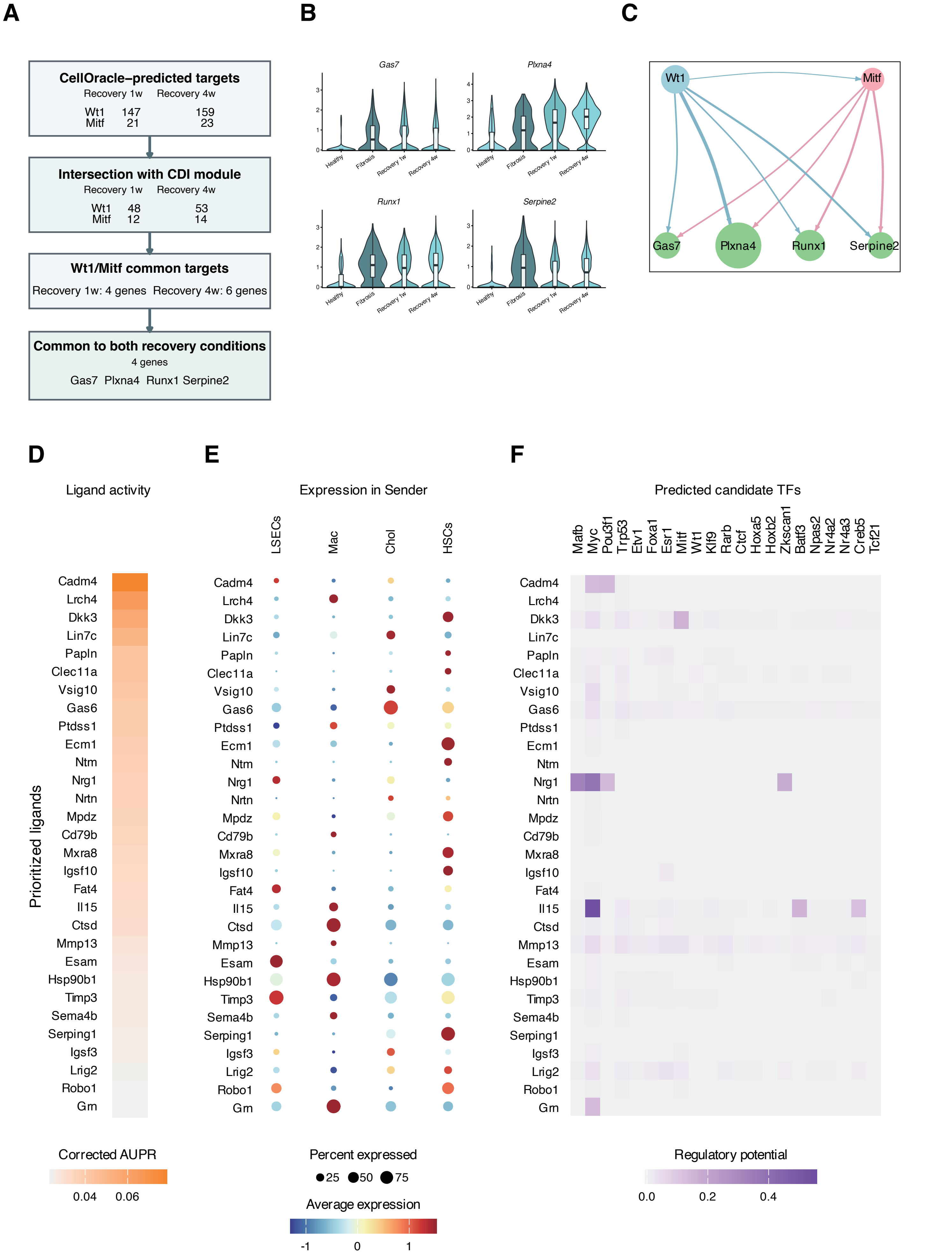

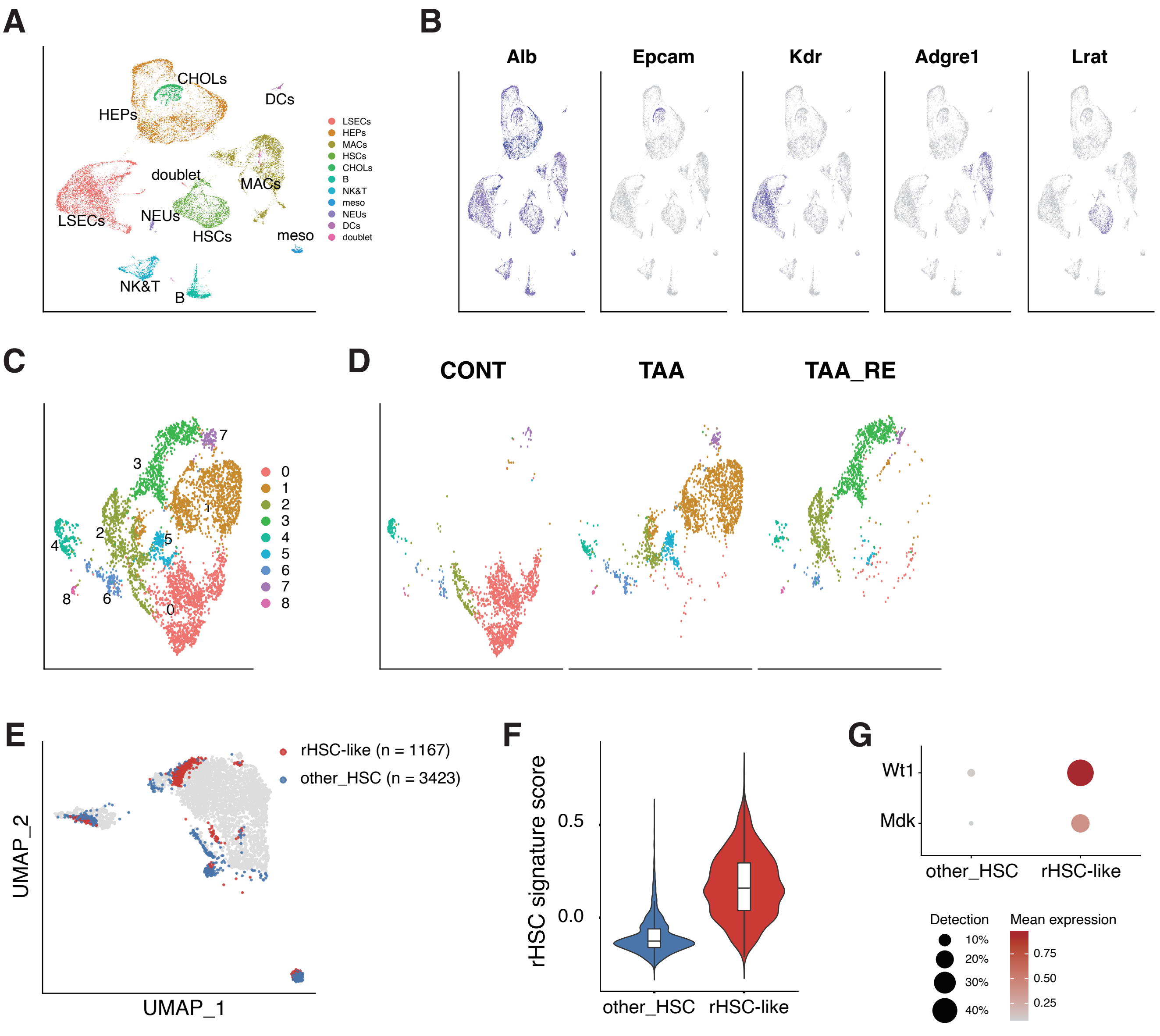


**H**

| condition | sample_id | total_cells | rHSC-like_cells | rHSC_mapping_proportion (%) |
|---|---|---|---|---|
| CONT | CONT_WT1198 | 446 | 48 | 10.8 |
| CONT | CONT_WT1199 | 484 | 30 | 6.2 |
| CONT | CONT_WT1200 | 553 | 39 | 7.1 |
| TAA | TAA_WT11 | 656 | 274 | 41.8 |
| TAA | TAA_WT13 | 558 | 157 | 28.1 |
| TAA | TAA_WT14 | 598 | 278 | 46.5 |
| TAA_RE | TAA_RE_WT1815 | 374 | 96 | 25.7 |
| TAA_RE | TAA_RE_WT1816 | 460 | 111 | 24.1 |
| TAA_RE | TAA_RE_WT1817 | 461 | 134 | 29.1 |

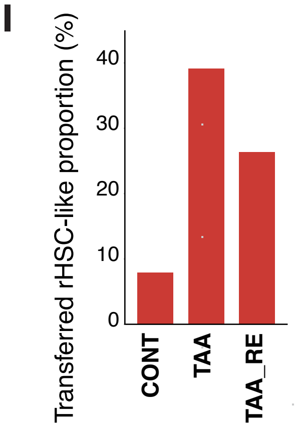


**J**

| metric | value |
|---|---|
| Shared reference variable features | 1,730 |
| Common RNA genes | 14,955 |
| Transfer anchors | 322 |
| Query cells | 4,590 |
| Reference cells | 3,771 |

Cross-species evaluation of the murine rHSC-associated transcriptional program in human liver datasets.

Supplementary Figure 7

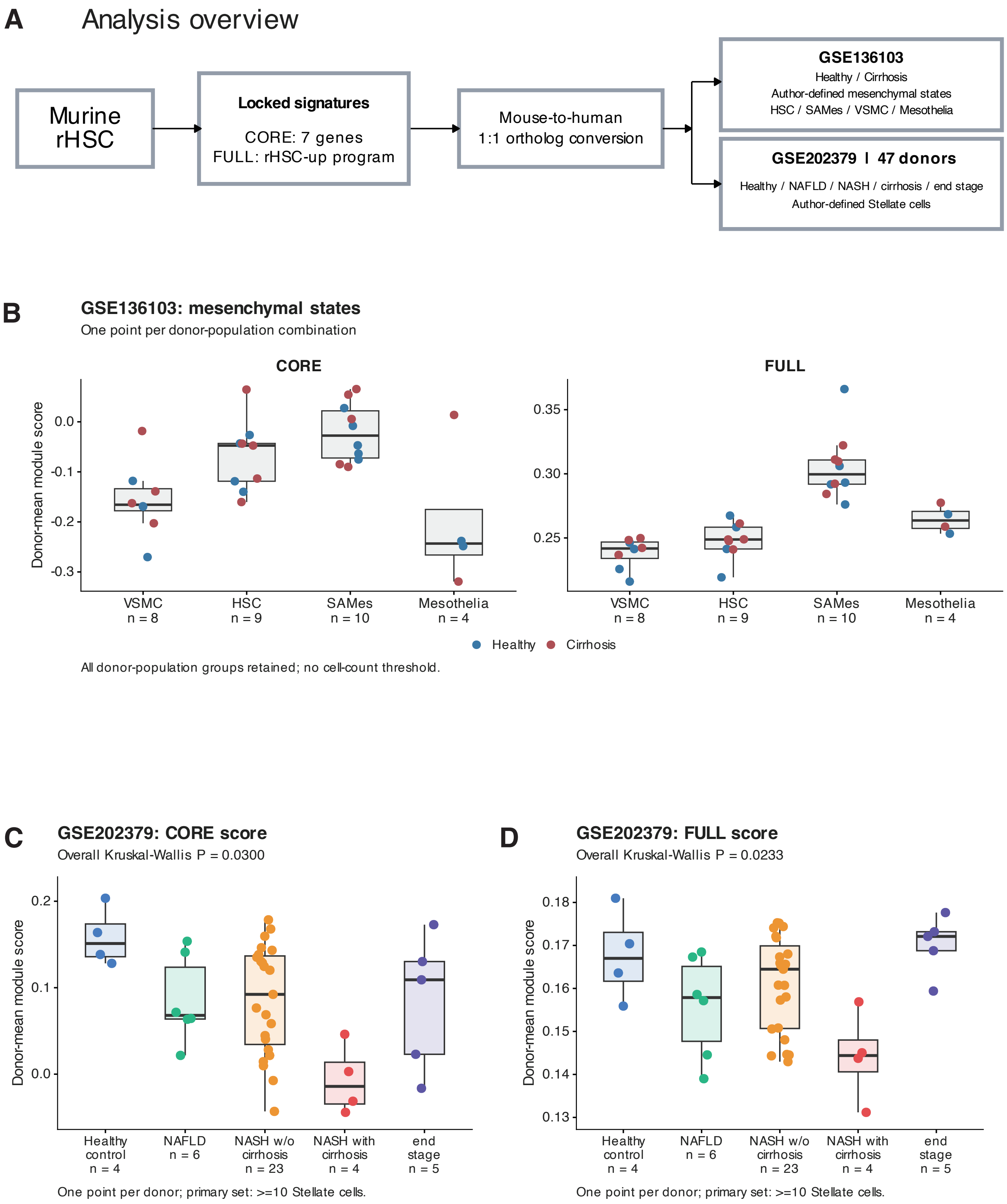

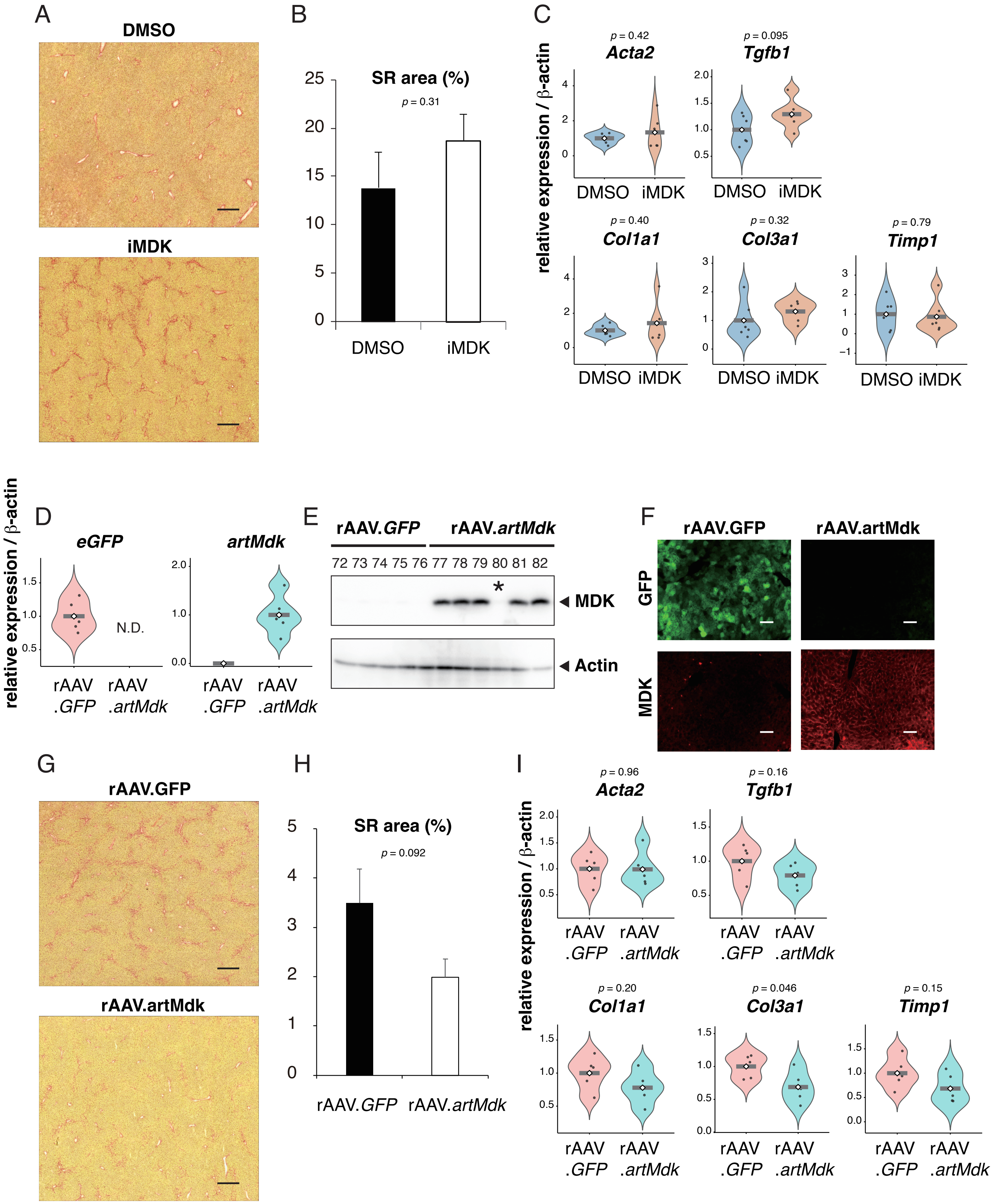
A
DMSO
iMDK
B
SR area (%)
p = 0.31
DMSO
iMDK
C
relative expression / β-actin
p = 0.42
Acta2
p = 0.095
Tgfb1
p = 0.40
Col1a1
p = 0.32
Col3a1
p = 0.79
Timp1
DMSO iMDK
D
relative expression / β-actin
eGFP
artMdk
N.D.
rAAV .GFP
rAAV .artMdk
E
rAAV.GFP
rAAV.artMdk
72 73 74 75 76 77 78 79 80 81 82
MDK
Actin
F
rAAV.GFP
rAAV.artMdk
GFP
MDK
G
rAAV.GFP
rAAV.artMdk
H
SR area (%)
p = 0.092
rAAV.GFP
rAAV.artMdk
I
relative expression / β-actin
p = 0.96
Acta2
p = 0.16
Tgfb1
p = 0.20
Col1a1
p = 0.046
Col3a1
p = 0.15
Timp1
